# Differential Chromatin Architecture and Risk Variants in Deep Layer Excitatory Neurons and Grey Matter Microglia Contribute to Major Depressive Disorder

**DOI:** 10.1101/2023.10.02.560567

**Authors:** Anjali Chawla, Doruk Cakmakci, Wenmin Zhang, Malosree Maitra, Reza Rahimian, Haruka Mitsuhashi, MA Davoli, Jenny Yang, Gary Gang Chen, Ryan Denniston, Deborah Mash, Naguib Mechawar, Matthew Suderman, Yue Li, Corina Nagy, Gustavo Turecki

## Abstract

Major depressive disorder (MDD) associated genetic variants reside primarily in the non-coding, regulatory genome. Here we investigate genome-wide regulatory differences and putative gene-regulatory effects of disease risk-variants by examining chromatin accessibility combined with single-cell gene-expression profiles in over 200,000 cells from the dorsolateral prefrontal cortex (DLPFC) of 84 individuals with MDD and neurotypical controls. MDD-associated accessibility alterations were prominent in deep-layer excitatory neurons characterized by transcription factor (TF) motif accessibility and binding of nuclear receptor (NR)4A2, an activity-dependent TF responsive to pathological stress. The same neurons were significantly enriched for MDD-associated genetic variation disrupting cis-regulatory sites and TF binding associated with genes involved in synaptic communication. Furthermore, a grey matter microglial cluster exhibited differentially closed chromatin in MDD affecting binding sites bound by TFs known to regulate immune homeostasis. In summary, our study points to specific cell types and regulatory mechanisms whereby genetic variation may increase predisposition to MDD.

## Introduction

Major depressive disorder (MDD) is a debilitating and life-threatening psychiatric disorder affecting almost 5% of the world’s population ^1,2^. Many neurobiological factors and cellular alterations have been previously associated with MDD including changes in monoaminergic and glutamatergic systems ^3–5^, and abnormalities in astrocytes ^6,7^, oligodendrocytes ^8^, and immune cells ^9,10^. In spite of progress, the precise molecular mechanisms mediating risk for MDD remain unknown.

Genome-wide associations studies (GWAS) have identified over 200 MDD risk loci ^11–13^; however, similarly to other psychiatric phenotypes, characterizing the functional impact of such variants has remained challenging as they mostly map outside protein-coding regions. In addition to genetic variation, the adverse effects of environmental factors, such as early-life stress, on increasing the risk for depression ^14,15^, are mediated by regulatory changes^16^. Single-cell atlases of open chromatin cis-regulatory elements (cCREs) holds promise to uncover mechanisms whereby disease-associated risk-variants may impact cell-specific gene-regulatory targets ^17,18^.

Previously, using single-nucleus RNA-seq (snRNA-seq), our group identified gene-expression changes mainly in the deep-layer excitatory neurons, and certain glial subtypes, including microglia in individuals with MDD ^19,20^. Here, we bridge current knowledge gaps by employing single-nucleus assay for transposase-accessible chromatin with sequencing (snATAC-seq)^21^, which provides unprecedented opportunities for interrogating epigenetic and genetic variants through the lens of chromatin accessibility ^17,18,22^. By measuring chromatin accessibility in more than 200,000 cells from the DLPFC of 84 individuals and integrating snRNA-seq obtained from the same samples ^19,20^, we generated a large cell-type specific accessibility and expression atlas of cortical cells in MDD.

Our results revealed that MDD-associated chromatin accessibility alterations were most prominent in NR4A2+ deep-layer excitatory neurons (ExN1) and in a grey matter (GM) microglial cluster (Mic2). ExN1 neurons were also most significantly enriched for the heritability of MDD GWAS SNPs. Examining allele-specific effects of MDD-associated genetic variation in these neurons identified TFs and genes likely affecting synaptic communication and plasticity. Together, our single-nucleus multi-modal dataset provides an in-depth examination of MDD-associated genetic variation and downstream gene regulatory mechanisms with cell type specificity.

## Results

### Chromatin accessibility in human dorsolateral pre-frontal cortical (DLPFC) cells

We performed snATAC-seq in 44 individuals who died during an episode of MDD and 40 age- and sex-matched neurotypical controls (Fig 1A, schematic; demographic and sample characteristics; Supplementary Table 1). Male and female nuclei were combined prior to microfluidic-capture, common variants, and chromatin accessibility measured in this dataset permitted accurate demultiplexing of pooled subjects (Supplementary Fig 1; Supplementary note). Filtering of doublets and low-quality barcodes revealed 201,456 high-quality nuclei having overall high TSS enrichment score (median: 7) and uniquely mapped fragments (median: 15,833) with equal contributions from both sexes (51% female cells) and conditions (53% case cells; Supplementary Figure 2-4).

**Figure 1.**
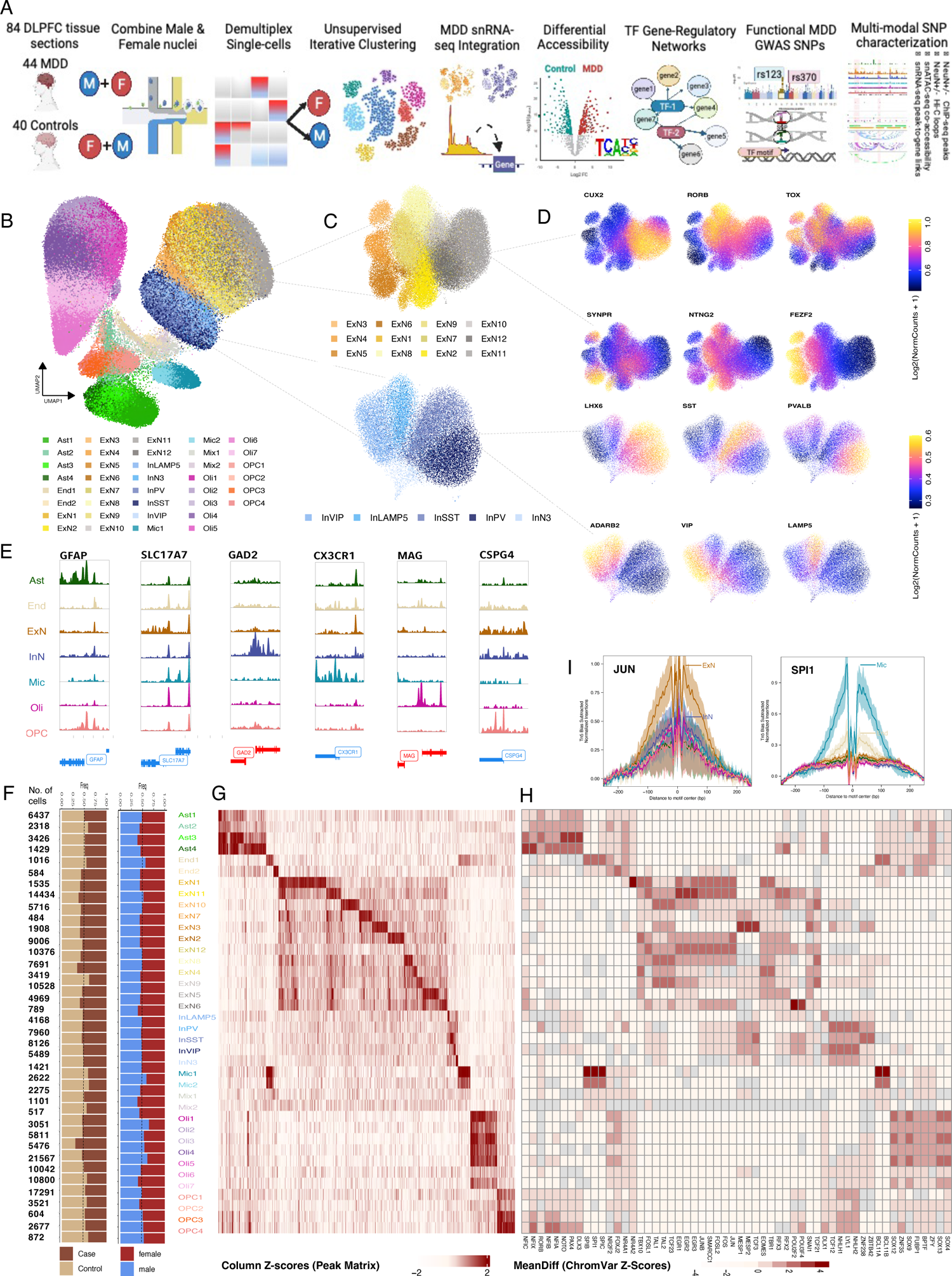
Chromatin Architecture at the single-cell level. A. Schematic overview of the snATAC-seq study design in 84 subjects from nuclei extraction and multiplexing to downstream data analyses B. UMAP of 201,456 Chromatin accessibility defined high-quality cells colored by 38 clusters from 7 broad cell-types. C. Iterative clustering within broad excitatory and inhibitory cell-types revealed 12 excitatory (ExN1-12; top), 5 inhibitory clusters (InN3/PV/SST/LAMP5/VIP; bottom). D. UMAP plot colored by gene-activity computed using ArchR GeneScore matrix for cortical layer (top) and interneuron lineage marker genes (bottom). Top: CUX2: Upper layer (L1-2), RORB, TOX: Middle layer (L3-4), SYNPR, NTNG2, FEZF2: Deep layer (L5-6). Bottom: LHX2, SST, PVALB: medial ganglionic eminence (MGE), ADARB2, VIP, LAMP5: caudal (CGE) lineage interneurons. E. Pseudo-bulk chromatin accessibility at cell-type marker genes (normalized by reads in TSS; Ast: GFAP; ExN1: SLC17A7; InN: GAD2; Mic: CX3CR1; Oli: MAG; OPC: CSPG4). F. Total no. of nuclei in each cluster followed by proportion of nuclei contributed by MDD and control subjects and contributed by male and female subjects. G. Heatmap showing differentially accessible cCREs identified in each cluster compared to all others (marker cCREs; Wilcoxon test, FDR < 0.05, Log2FC > 0.5). H. Heatmap showing mean differences in ChromVar based bias-corrected motif accessibility deviation z-scores in each cluster compared to all others (Top 3 marker TFs are plotted; Wilcoxon test, FDR < 0.05, MeanDiff>1.5; Grey boxes represent NA values). I. Tn5 bias subtracted motif footprints for cell-type marker TFs (JUN: Neuronal marker, SPI1: Microglial marker).

Unsupervised iterative clustering of nuclei by genome-wide chromatin accessibility measured at 500bp resolution ^23^ revealed 7 broad cell-types (ExN: 35.2%, InN: 13.4%, Ast: 6.8%, Oli: 36.8%, OPC: 3.8%, Mic: 2.4%, End: 1.6%) consisting of 38 distinct clusters (Fig 1B; Extended Fig 1D). Multiple approaches for assigning cell annotations confirmed snATAC-seq cell-type and cluster identities (Fig 1C-E, Extended Fig 1-2), including label-transfer from sample-matched snRNA-seq ^20^ (median: 0.94; Extended Fig 1A-E) and published PFC snATAC-seq ^24^ (median: 0.99; Extended Fig 2A-B).

**Figure 2.**
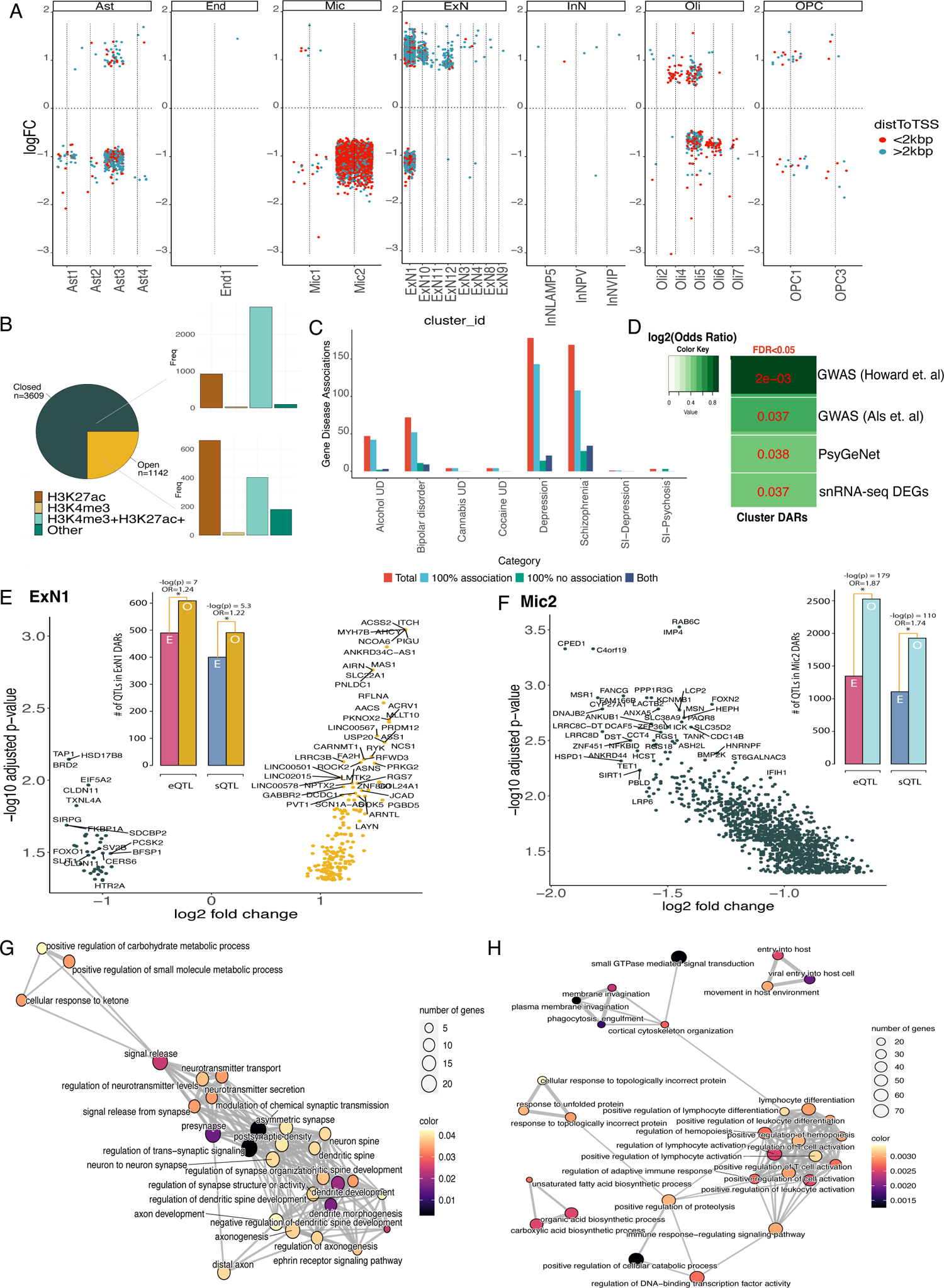
Chromatin profiling in individuals with major depression. A. Differentially accessible regions (DARs) in MDD compared to neurotypical control by cluster (Limma-voom; FDR < 0.05 & abs(LogFC) > log2(1.1)). Points above the line represent increased accessibility, below decreased. Colors represent distance to the nearest transcription start sites (TSS). B. Total number of differentially accessible regions (DARs) that are differentially more (open; n=1142) or less accessible (closed; n=3609) in MDD versus controls subjects across all clusters. Adjacent bar plots show the total number of PsychEncode ChIP-seq histone modification peaks from cortical cells (H3K27ac; marks active enhancers and H3K27ac+H3K4me3; marks promoters) that overlap differentially more (bottom) or less (top) accessible MDD DARs. C. PsyGeNet identified the total number of gene-disease associations linked to DARs across all clusters (DAR-linked genes (r>0.45); SI: substance induced; 100% association: consistent evidence for the disease; 100% no association: consistently not associated with the disease). D. Gene overlap enrichment (one-sided Fisher’s exact test; FDR<0.05) between DAR-linked genes and MDD-associated GWAS risk-genes from Howard et. al (2019) and Als et.al (2023), PsyGeNET genes associated with depressive disorder, and MDD DEGs identified in snRNA-seq meta-analysis across both sexes. E. DAR-linked genes plotted according to log2FC and -log10(adjusted p-values) of each DAR in ExN1 (left) and F. Mic2 (right). Inset: Bar plots depict significant enrichment (p<0.05) of expression (eQTLs) and splicing quantitative trait loci (sQTLs) in ExN1 (left) and Mic2 DARs (right) compared to expected genomic coverage. G-H. Gene ontology (GO) terms enriched in G. ExN1 and H. Mic2 DAR-linked genes converted into networks (emapplot) plotted by significance (FDR<0.05). Nodes in the network show significant GO terms (size of node is scaled by the number of genes associated with each GO term). Edges connect overlapping gene sets (minimum 10% overlap required for an edge criterion between GO terms and thickness of edges are scaled by the overlap percentage).

The clusters were characterized by cluster-specific accessibility across 239,824 cCREs, (hereafter called marker cCREs; Fig 1G; Wilcoxon test, FDR<0.05; Extended Fig 1I (top); Supplementary Table 2) and cluster-specific TF motif accessibility and binding (Wilcoxon test, FDR<0.05, Fig 1H-I, Supplementary Table 3). To identify gene-regulatory mechanisms in cortical cells, we examined correlations between cCRE accessibility and gene-expression (peak-to-gene linkages), identifying 117,328 unique cCREs highly correlated (r>0.45) with 11,575 expressed genes (Supplementary Table 4). As expected, almost all (>90%) of these cCREs overlapped histone markers of transcriptionally active chromatin from cortical cells ^25^ (Extended Fig 1I (bottom)) and captured gene-regulatory variations from each cell-type (Extended Fig 1C).

### MDD-associated chromatin differences found in microglia and deep-layer neurons

We investigated MDD-associated accessibility within clusters and cell-types using per-subject pseudo-bulked accessibility estimates (Method: Differential accessibility analysis; Supplementary Table 5). Of 38 characterized clusters, 25 clusters were associated with a total of 4751 MDD-associated differentially accessible regions (hereafter called DARs; Fig 2A), whereas only 297 differential chromatin regions were found at broad level (Supplementary figure 5A-C). Most DARs were less accessible in individuals with MDD (76%; Fig 2B) and were primarily observed in clusters of microglia Mic2 (58.5%) and excitatory neurons ExN1 (21.2%). The number of cells in these clusters were comparable to all others (except Oli4; interquartile range (IQR)) and depicted uniform contributions from potential covariates (Fig 1F; Supplementary Fig 4).

To understand the likely regulatory impact of DARs, we asked how they overlapped with PsychEncode ChIP-seq histone marks from cortical cells ^25^. Nearly all of the DARs less accessible in MDD (98%) coincided with histone marks that are typically found in gene promoters (defined by H3K4me3+H3K27ac; 74%) and enhancers (H3K27ac, 23%). By contrast, most DARs more accessible in MDD (85%) overlapped with enhancers (H3K27ac; 56%) more than promoters (28%; Fig 2B).

To understand the overall functional impact of DARs, we identified the most likely affected genes (n= 2254) using peak-to-gene linkages (r>0.45) and investigated their known functions. Of these genes, the highest psychiatric disease association was found for depression (Fig 2C), whereby 80% of these associations showed consistent evidence in PsyGeNet ^26^ and 64% for schizophrenia. In fact, there was an overrepresentation among DAR-linked genes (FDR < 0.05; one-sided Fisher’s exact test; Fig 2D) for PsyGeNet genes linked to depression, MDD-associated genetic variation ^11,13^ and genes previously observed to be differentially expressed in MDD ^20^. We did not observe similar enrichments for genes associated with cell-type DARs observed at broad level.

Cell type-specific roles of proximal and distal gene-regulation have been previously reported. For example, activity-dependent regulation in neurons involves many distal regulatory elements ^27^ while microglial-activation involves mostly proximal regulation ^28^. Here we similarly observed that whereas DARs in excitatory neurons tended to be distal to any TSS (e.g. 77% of DARs in ExN1 were > 2kbp from a TSS; Fig 2A), DARs in microglia tended to be proximal to a TSS (e.g. 73% of DARs in Mic2 were within 2kbp of a TSS).

Given that ExN1 and Mic2 were the most affected clusters in MDD, we focused on these to examine the relationship between altered chromatin and variants associated with gene regulation. Both ExN1 and Mic2 DARs showed significant enrichment for splicing (sQTLs) and expression quantitative trait loci (eQTLs; Fig2E-F inset), implicating potential impact of DARs on DLPFC gene expression. Moreover, using snRNA-seq clusters^20^, we identified significantly increased module scores in MDD for expression of genes linked to ExN1 DARs showing specifically increased accessibility in MDD (p = 0.0009, Wilcoxon rank-sum test; Extended figure 2F), and we observed tendency for the opposite in Mic2 (p=0.1).

Finally, to elucidate functional pathways likely affected by MDD-associated chromatin, we examined cluster-specific DAR-linked gene ontologies (GO; Supplementary Table 6). ExN1 DARs were largely more accessible in cases (73% of DARs) and linked to genes involved in modulating synaptic and neurotransmitter release functions (FDR<0.05; Fig 2H). Whereas Mic2 DARs with significantly decreased accessibility in MDD linked to genes enriched in phagocytosis- and immune-signaling pathways (FDR<0.05; Fig 2I; Supplementary figure 5D).

### TF binding sites overrepresented in MDD-associated chromatin

TF binding can both regulate and be affected by chromatin accessibility leading to gene-expression ^29^. In order to determine the possible interplay between TFs and transcriptional effects, we first identified TF binding motifs enriched in cluster-specific DARs compared to GC-matched background peaks (Homer ^30^; p<0.05, Fig 3A, Supplementary table 7). TF binding motifs highly enriched (FDR<0.05) in ExN1 DARs belonged to basic leucine zipper (bZIP), basic helix–loop–helix (bHLH), Zinc finger (Zf), and helix-turn-helix (HTH) families, which are known to regulate activity-dependent ^31^ and neurodevelopment-associated functions ^32,33^.

**Figure 3:**
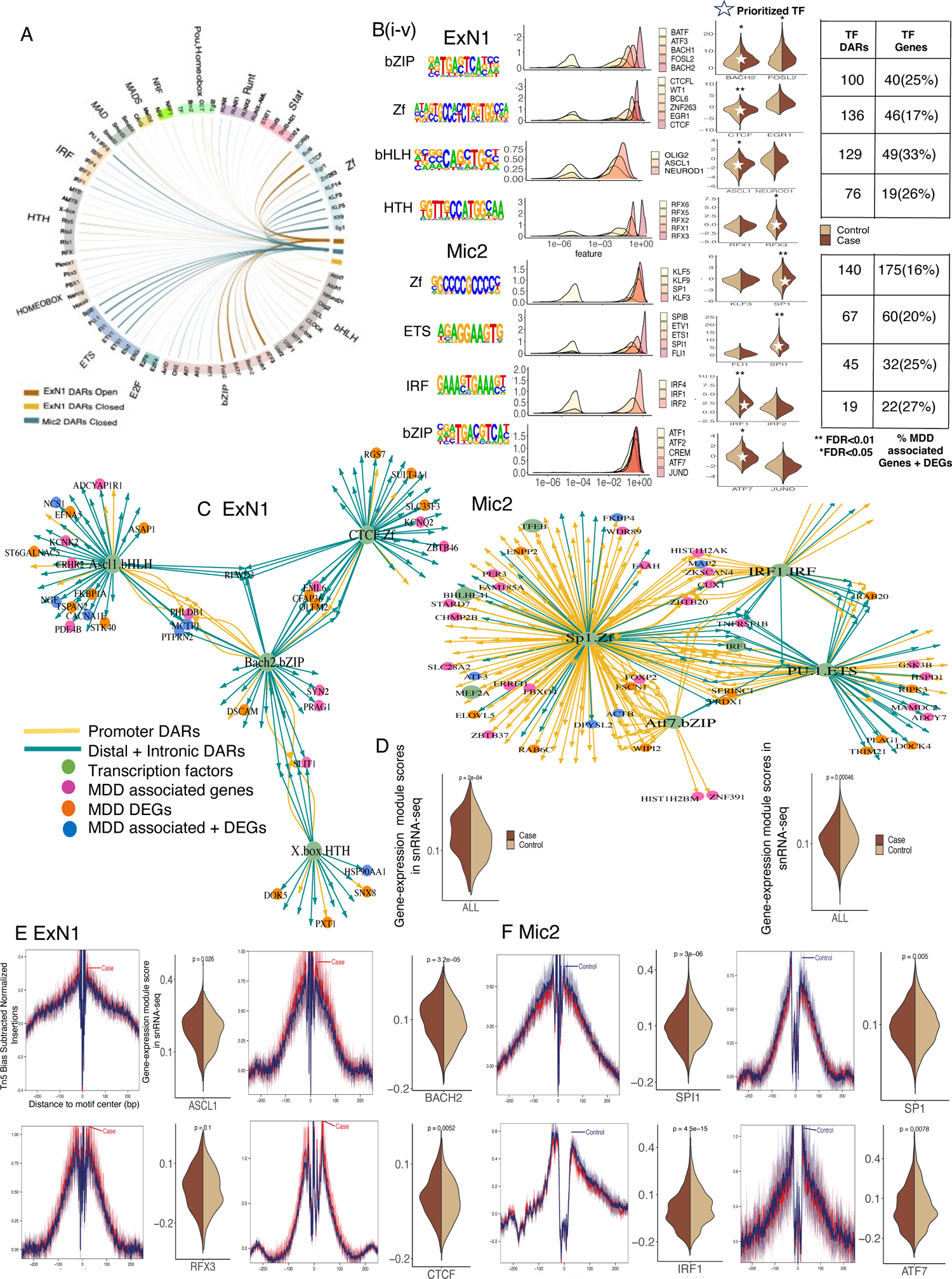
The role of transcription factors in MDD. A. Circos plot shows TF motif families enriched (Homer; p<0.05) in differentially more (open) or less accessible (closed) ExN1 DARs and differentially less accessible (closed) Mic2 DARs (width of links represent –log10(p-value) scaled within each cluster). Each segment is colored by individual TF families representing TF motifs significantly enriched in DARs. Only TFs present in both ArchR peak-motif matrix and snRNA-seq integrated gene-expression matrix are plotted. B: Workflow for TF prioritization. Left to right: (i) Top 4 most significant TF families (FDR<0.05) in ExN1 (top) and Mic2 (bottom); (ii) DNA motifs of prioritized TFs in each TF family; (iii) Density plots show snRNA-seq integrated gene expression for topmost significant TFs belonging to each family (plotted from the lowest to highest median gene expression) across cells in each cluster (iv) ChromVar based bias-corrected TF motif accessibility deviation z-scores in MDD versus controls (Wilcoxon test, **FDR<0.01, *FDR<0.05; white stars indicate prioritized TFs with the most significant differences in motif accessibility z-scores in MDD versus controls; (v) Number of DARs enriched for the prioritized TF motifs and the number of target-genes associated with prioritized TFs (% represents TF target-genes which are also MDD DEGs (in MDD bulk or single-nucleus RNA-seq datasets) or MDD-associated genes (in MDD GWAS or PsyGeNET depressive disorder genes). C. TF target-gene regulatory networks for prioritized TFs associated with ExN1 DARs (FDR<0.05; left) and Mic2 DARs (FDR<0.01; right). Nodes in the network are colored by target-gene status of MDD associated genes or MDD DEGs or both. Edges are colored by type of DARs (promoters, introns or distal DARs). An edge is added to the network when a significant DAR contains the prioritized TF motif and connects to genes linked to any DAR through peak-to-gene linkages (r>0.45) and nearest genes to promoter DARs. D. snRNA-seq gene expression module scores for the prioritized TF target-genes in ExN1 (left) and Mic2 (right) compared between MDD versus control groups (Wilcoxon test) in the corresponding snRNA-seq clusters. E-F. Tn5 bias subtracted motif footprints of prioritized TFs in MDD versus control subjects (left) and snRNA-seq gene-expression module scores for target-genes of prioritized TFs in MDD versus controls in corresponding snRNA-seq clusters (right; Wilcoxon test) for E. ExN1 and F. Mic2.

On the other hand, TF motifs enriched (FDR<0.05) in Mic2 DARs included many canonical microglial factors, such as ETS domain (SPI1) and interferon regulatory factors (IRFs), some of which are pioneering and lineage-specifying TFs that can regulate local chromatin accessibility ^34,35^ and can affect microglial development ^36^, proliferation ^37^, and immune-activation^34,38^.

To confirm these observations, we assessed GC and accessibility bias-corrected TF motif deviations across motif-matching cCREs in each cluster (ChromVar ^39^; Wilcoxon test; FDR<0.05). These results suggested an overall agreement between MDD-associated TF motifs enriched in differentially more or less accessible DARs among MDD individuals and those identified comparing motif accessibility signals between MDD and controls at single-cell resolution (Supplementary Table 8).

### TFs mediate effects of MDD-associated chromatin on gene expression

We hypothesized that reduced chromatin accessibility in MDD at TF binding sites would reduce TF binding and result in reduced expression of target genes, and vice versa. We investigated this hypothesis by identifying a set of TFs whose binding was most likely to be affected by MDD-associated accessibility, compiling a list of their most likely target genes, and then examining if the expression of these genes was associated with MDD.

To identify high-priority TFs affected by differential accessibility, we selected TFs with motifs enriched in DARs that were also highly expressed in integrated snRNA-seq. Then, among these, we further restricted selection to those with the greatest MDD-associated accessibility differences at their binding sites in cases versus controls (Fig 3B; Supplementary Figure 5E).

We identified the likely targets of these prioritized TFs by constructing gene regulatory networks using igraph (Fig 3C; Method: Gene regulatory networks) in ExN1 (BACH2, ASCL1, CTCF, RFX3) and Mic2 (SP1, SPI1, IRF1, ATF7). The nodes in green represent non-coding DARs with individual TF binding motifs connected to their putative target-genes colored according to MDD DEGs or MDD-associated genetic variation and PsyGeNET depressive disorder gene status.

Finally, we calculated MDD-associated module scores for these networks using snRNA-seq clusters^20^. In general, these scores confirmed that increased binding site accessibility was associated with increased module expression levels, and vice versa (Fig 3D-F).

### Strong enrichment for MDD heritability in deep-layer excitatory neurons

To prioritize cell type clusters that are likely to mediate the heritable risk for MDD, we calculated GWAS SNP heritability enrichment within cluster-specific chromatin using stratified linkage-disequilibrium score regression (S-LDSC; Method: Heritability enrichment) ^40,41^. For this, we used GWAS summary statistics available for MDD and related psychiatric phenotypes and non-brain related control traits. The heritable risk for psychiatric disorders and traits that are known to be genetically correlated with MDD ^42^ were significantly enriched in marker cCREs of ExN1 and ExN2 clusters (Fig 4A). Similarly to ExN1, ExN2 showed similarities to deep-layer (layer 5-6) excitatory neurons (Extended Fig 2A, 3C). In fact, enrichments calculated from data obtained from both MDD GWAS used in our study^11,13^ were coherently highest in ExN1 cluster (Fig 4A; p=0.003 (Howard), p=0.0005 (Als), Supplementary Table 9). We also calculated heritability enrichments in MDD-associated chromatin in each cluster (candidate DARs), showing that the enrichment was likewise highest in ExN1 for MDD (Fig 4B; p=0.01; nominal) and insomnia (p=0.02).

**Figure 4.**
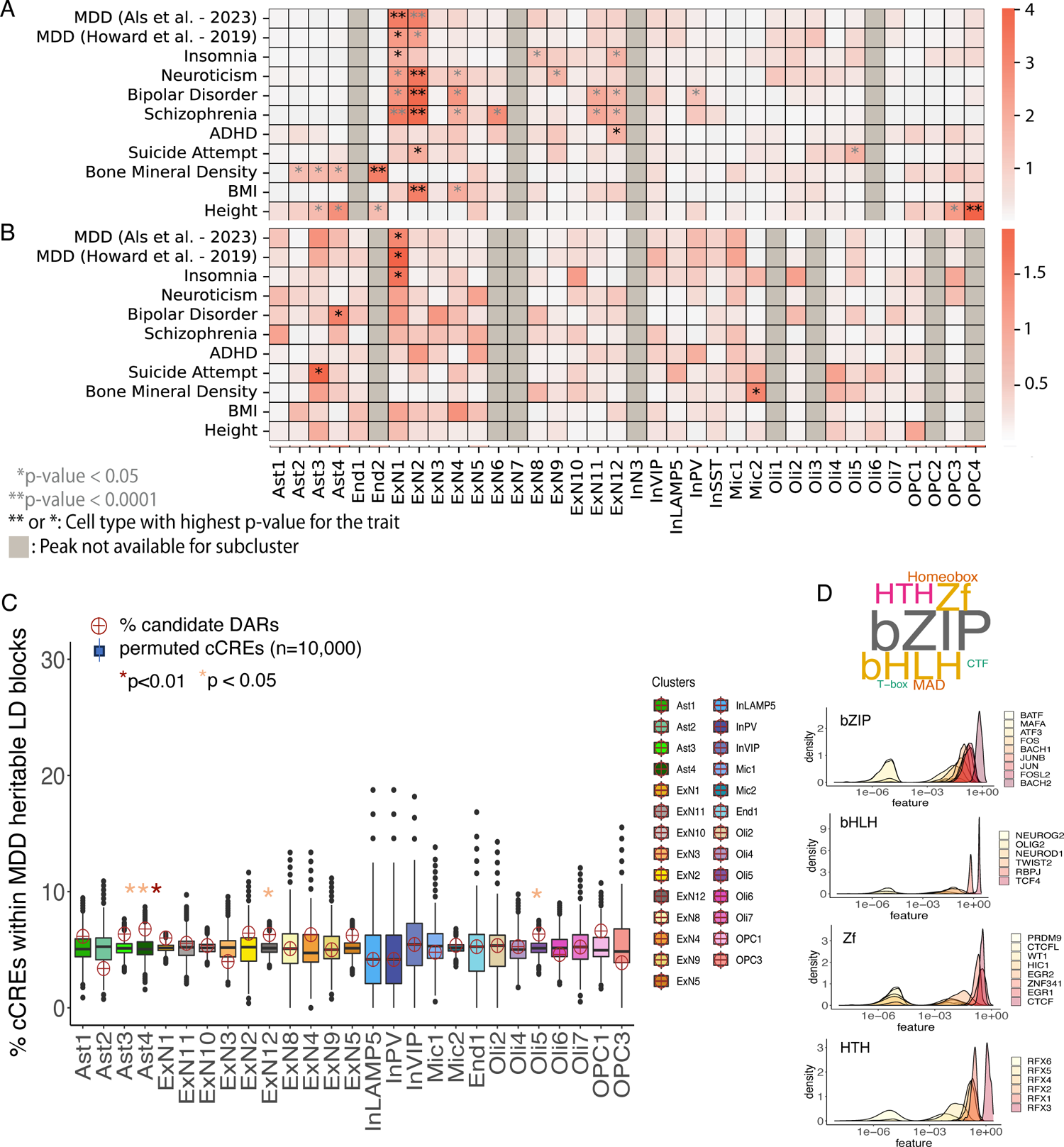
Genetic overlap with cell- and MDD-specific chromatin structure. A-B: Heatmaps scaled by –log10(p-value) (LDSC z-score test) for heritability enrichment of GWAS SNPs belonging to each trait in each cluster using A. cluster-specific marker cCREs B. cluster-specific candidate DARs C. Overrepresentation of cluster-specific candidate DARs (permutation test; n=10,000; p<0.05) within significant heritable LD blocks identified for MDD (HESS; p<0.05). Boxplots represent expected proportions of chromosome-matched permuted cCREs within any LD block. Circles show the observed proportion of candidate DARs within significantly heritable MDD-associated LD blocks. D. Word cloud represents TF motif families significantly enriched (q<0.05; Homer) in both ExN1-specific marker cCREs and candidate DARs located within MDD-associated LD blocks. The size of TF families is scaled by the total number of distinct TF members (belonging to these families) identified to be significantly enriched (q<0.05). Density plots shows snRNA-seq integrated gene-expression of enriched TFs across ExN1 cells.

To determine if these enrichments were driven by linkage disequilibrium (LD), we first estimated the heritability of MDD-associated variants in heritable LD blocks defined by 1000 Genome European (EUR) population using heritability estimator from summary statistics (HESS) ^43^. This resulted in about 5% of LD blocks significantly enriched for MDD-associated genetic variants (p<0.05, Supplementary table 10). We then examined if MDD-associated blocks contained an overrepresentation of candidate DARs within each cluster compared to chromosome-matched cCREs in any LD block (Permutation testing; nPerm=10,000). Consistent with previous observations, the most significant enrichment was observed in ExN1 (p=0.007; Fig 4C).

Finally, to identify regulatory TFs associated with chromatin enriched for MDD heritability, we examined TF motifs enriched in ExN1-specific cCREs located within MDD-associated LD blocks compared to GC-matched background peaks. TF families that were significantly enriched (q<0.05; Supplementary table 11) in both ExN1 marker and differential chromatin included those associated with activity-dependent functions (bZIP; Fig 4D) ^31^ and neurodevelopment (bHLH, HTH, Zf) ^32,33^.

### MDD-associated genetic variation shows cell type specific effects on gene regulatory sites

To examine the functional impact of MDD-associated genetic variation within each cluster, we assessed allelic-effects of MDD-associated SNPs on accessibility of cCREs and TF binding sites and identified likely target risk genes. We used MDD-associated SNPs in GWAS ^11–13^, those in high LD with them (r2 value ≥0.8) and those identified by finemapping ^11,44^. To predict accessibility in each cluster, we used three complementary models; in-silico mutagenesis (ISM), delta-support vector machine (delta-SVM) ^45^ and gapped-kmer (gkm-explain) ^46^ based variant-effect scores derived from cluster-specific gkmSVM classifiers ^47,48^ (Supplementary Fig 6-7).

In order to determine cell-type and cluster-specific effects of MDD-associated genetic variation, we focused on MDD-associated SNPs overlapping cell-type- or cluster-specific candidate cCREs (referred to as candidate SNPs (cdSNPs) with 14% cdSNPs in candidate DARs and remaining in marker cCREs; Extended Fig 3A-B; Supplementary Table 12). Compared to MDD-associated SNPs, cdSNPs showed significant enrichment for expression quantitative trait loci (eQTLs; p<1×10^-3^; Hypergeometric test) and splicing QTLs (sQTLs; p<5×10^-8^), suggesting an over-representation of MDD SNPs with known effects on DLPFC gene-expression using our candidate cCREs (Extended Fig 3C-D).

Overall, we identified 97 non-coding statistically significant cdSNPs (sSNPs; 15% in candidate DARs; Extended Fig 3B) that were significant across all three gkmSVM models. Notably, 97.9% of MDD sSNPs were exclusively identified in clusters of a single broad cell-type (Extended Fig 3E), suggesting largely cell-type specific impact of genetic variation on chromatin accessibility.

Further, GWAS SNPs often alter expression of distally-located genes ^17^. Putatively long-range gene-regulatory effect of genetic variation was determined by identifying gene-expression targets most significantly correlated (up to 500kbps; peak-to-gene linkages) with accessibility of cCREs associated with MDD sSNPs. Both nearest and linked target genes of MDD sSNPs enriched for pathways related to synapse organization and communication (Extended Fig 4A-B; Supplementary Table 13).

### Functional impact of MDD-associated genetic variation in ExN1 and Mic2 clusters

To examine functional effects of MDD sSNPs in clusters that were strongly associated with MDD, we focused on sSNPs showing significant allelic-impact in ExN1 and Mic2 and overlapping cell-type- or cluster-specific candidate cCREs. We highlighted sSNPs that strongly disrupted accessibility in these cluster (medium- to high-confidence) and integrated PsychEncode DLPFC histone modifications ^25^ and 3D chromatin loops ^49^ to interpret long-range gene-regulatory targets (Method: Multi-modal Visualization).

Of all MDD sSNPs, 25% (52% s/eQTLs; Fig 5A) mapped to ExN1 marker cCREs or candidate DARs. Together, the target genes of ExN1 sSNPs (nearest, linked, and eQTLs) enriched for pathways related to synaptic communication and plasticity (Extended Fig 4C). This included a lead MDD sSNPs (rs2276138; Fig 5B) in GWAS meta-analysis ^50^ overlapping ExN1 marker cCRE in intronic region of *SPTBN2* gene and disrupting TF binding site for RFX3, whose motif accessibility was also altered in ExN1 DARs and at single-cell level in MDD (Fig 3B). Moreover, RFX3 motifs enriched in ExN1 cCREs located within heritable LD regions for MDD (Fig 4D) and have been previously associated with neuropsychiatric disorders ^33,51^. This sSNP is an eQTL and sQTL for a few genes (Fig 5A) including a GTEx eQTL, *ZDHHC24,* with major regulatory effects on synaptic plasticity ^52^ and associated through peak-to-gene linkages (r=0.42) with expression of *SYT12,* a synaptic vesicle protein. Additionally, MDD sSNP in *SHANK2* gene (rs11601579; Fig 5C) disrupted binding site accessibility for GLI2 TF, associated with neurodevelopment ^53^. This sSNP overlapped ExN1 marker cCRE and significantly correlated with the anti-sense RNA expression of *SHANK2* (r=0.6), a postsynaptic density scaffolding protein linked to neuropsychiatric disorders ^54,55^. Finally, about 40% (10/25) of ExN1 sSNPs were found to alter accessibility exclusively in this cluster (Extended Fig 5A-B).

**Figure 5:**
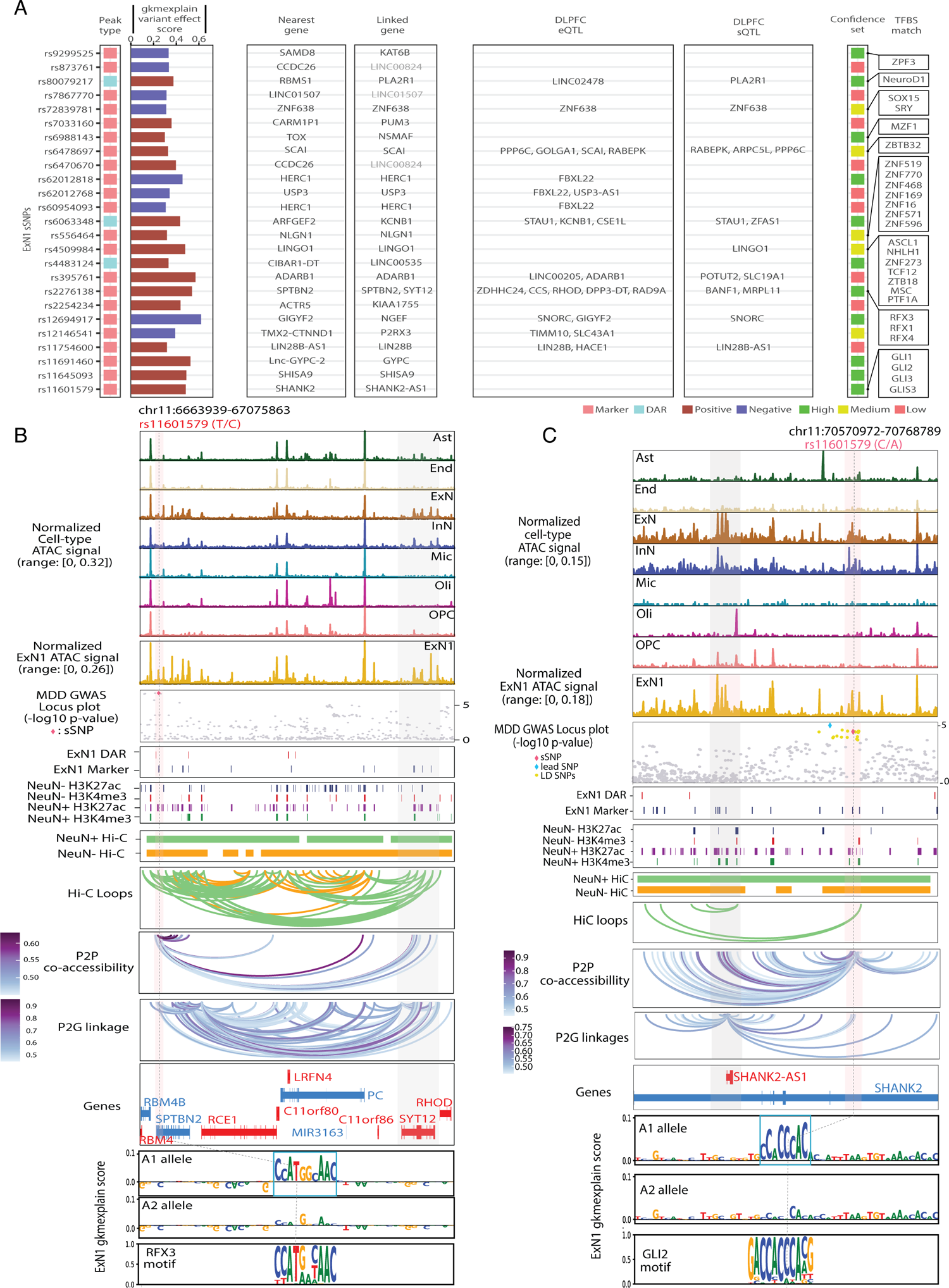
MDD sSNPs with significant allelic-effects on chromatin accessibility landscape in ExN1 cluster. A. Left to right: MDD sSNPs with significant allelic-effects in ExN1 cluster; ExN1 sSNP-associated nearest genes, linked genes most significantly (FDR<0.05) correlated with cCREs associated with sSNPs (peak-to-gene linkages; in grey are genes that did not pass FDR); DLPFC eQTLs and sQTLs genes associated with sSNPs; sSNPs categorized based on accessibility disruption confidence sets; TF binding sites potentially disrupted by sSNPs (listed in the order of significance for TF motif match). B. Example plot for ExN1 sSNP (rs2276138). Top to bottom: snATAC-seq-derived pseudo-bulk tracks for each cell-type and ExN1 cluster, Manhattan plot showing MDD sSNP (chr11: T>C; rs2276138) overlapping ExN marker cCRE (peak-to-gene linkage: SPTBN2 gene) and ExN1 marker cCRE (peak-to-gene linkage: SYT12 gene), ExN1 candidate DARs and marker cCREs within the plotted region, DLPFC neuronal and non-neuronal (NeuN+/-) H3K27ac and H3K4me3 ChIP-seq peaks merged across each subject 25, DLPFC NeuN+/- promoter anchored HiC fragments and HiC loops49, snATAC-seq peak-to-peak co-accessible loops (r>0.45) within 10kbps of sSNP associated cCRE, SYT12 gene locus (chr11:67006771-67050863) with the most significant peak-to-gene linkage (r=0.42) with cCRE associated with sSNP, gkmExplain importance scores for each base in 50-bp region surrounding sSNP for A1 and A2 alleles from ExN1 gkm-SVM model, the predicted TF binding site (RFX3) affected by sSNP is shown at the bottom. C. Example plot for ExN1 sSNP (rs11601579). Top to bottom: snATAC-seq-derived pseudo-bulk tracks for each cell-type and ExN1 cluster (normalized by reads in TSS), Manhattan plot showing MDD lead SNP, SNPs in LD (r>0.8), and MDD sSNP (chr11: C>A; rs11601579) overlapping ExN and ExN1 marker cCRE (peak-to-gene linkage: SHANK2-AS1 gene), ExN1 candidate DARs and marker cCREs within the plotted region, DLPFC neuronal and non-neuronal (NeuN+/-) H3K27ac and H3K4me3 ChIP-seq peaks merged across each subject25, DLPFC NeuN+/- promoter anchored HiC fragments and HiC loops49, snATAC-seq peak-to-peak co-accessible loops (r>0.45) within 10kbps of sSNP associated cCRE, SHANK2-AS1 gene locus (chr11-70625973-70635733) with the most significant peak-to-gene linkage (r=0.6) with cCRE associated with sSNP, gkmExplain importance scores for each base in 50-bp region surrounding sSNP for A1 and A2 alleles from ExN1 gkm-SVM model, the predicted TF binding site (GLI2) affected by the sSNP is shown at the bottom.

In Mic2, MDD-associated variants with allelic-effects on chromatin accessibility (Fig 6A) included a sSNP mapping to microglia cell-type marker cCRE (rs5995992; Fig 6B) and showing Hi-C linkages to *ARL4C* gene, which regulates lipid homeostasis in immune cells ^56^. This sSNP disrupted binding site for FLI1 TF, which is known to be involved in microglia development and homeostasis ^36,57^ and whose motifs were enriched in Mic2 DARs (Fig 3A). Additionally, a lead MDD GWAS SNP ^11,12^, which is an e/sQTL for several genes (rs5995992; Fig 6C) mapped to candidate Mic2 DAR in promoter region of *EP300,* a gene encoding histone acetyl-transferases and is co-activator of NF-κB mediated immune-activation ^58^.

**Figure 6:**
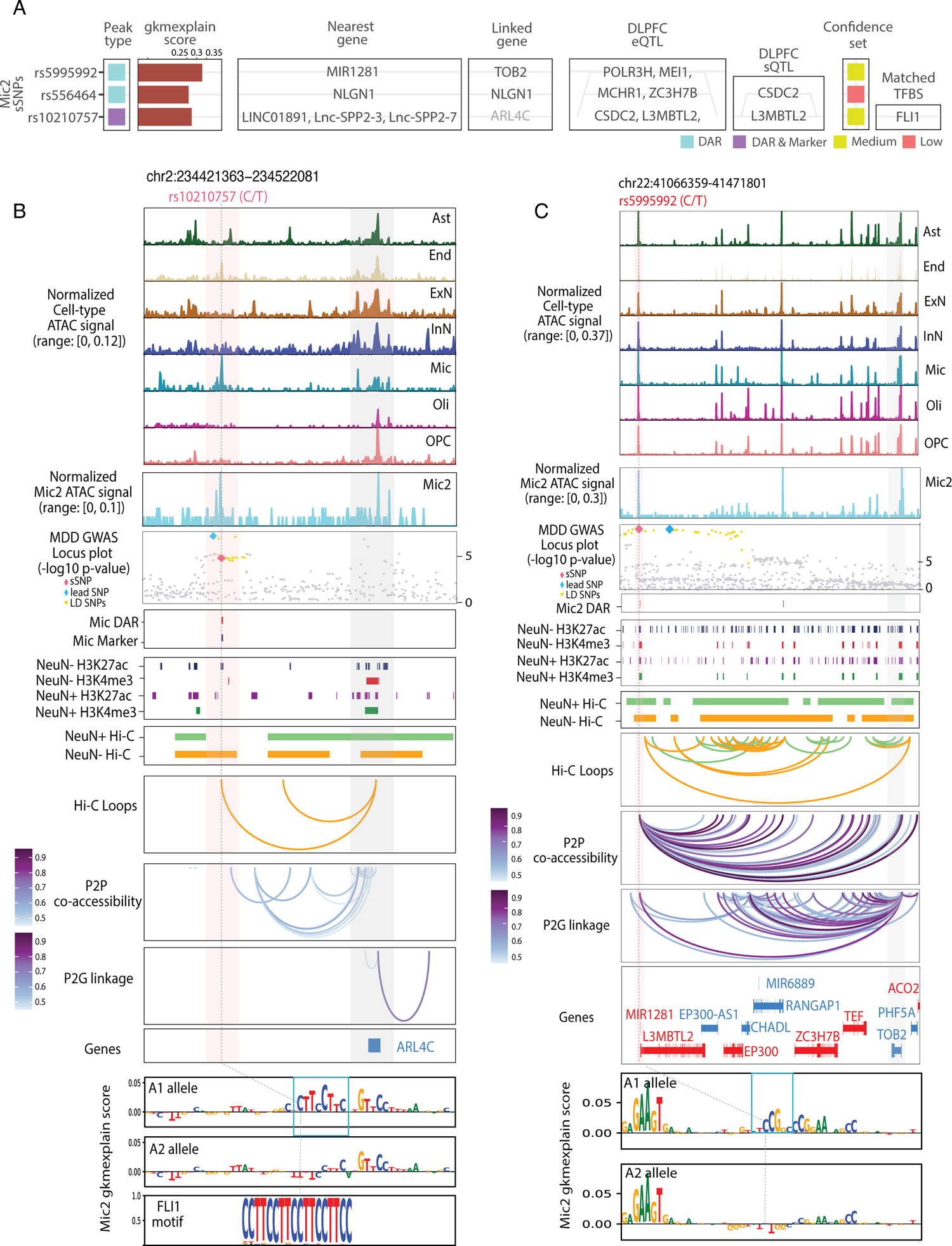
MDD sSNPs with significant allelic-effects on chromatin accessibility landscape in Mic2 cluster. A. Left to right: MDD sSNPs with significant allelic-effects in Mic2 cluster; nearest genes to sSNPs, linked genes most significantly (FDR<0.05) correlated with cCREs associated with sSNPs (peak-to-gene linkages, in grey are genes that did not pass FDR); DLPFC eQTLs and sQTLs genes associated with sSNPs; sSNPs categorized based on accessibility disruption confidence sets; TF binding sites potentially disrupted by sSNPs (listed in the order of significance for TF motif match) B. Example plot for Mic2 sSNP (rs10210757). Top to bottom: snATAC-seq-derived pseudo-bulk tracks for each cell-type and Mic2 cluster (normalized by reads in TSS), Manhattan plot showing MDD lead SNP, SNPs in LD (r>0.8), and MDD sSNP (chr2: C>T; rs10210757) overlapping Mic marker cCRE (also a candidate DAR in Mic; FDR=0.1), Mic candidate DARs and marker cCREs within the plotted region, DLPFC neuronal and non-neuronal (NeuN+/-) H3K27ac and H3K4me3 ChIP-seq peaks merged across each subject25, DLPFC NeuN+/- promoter anchored HiC fragments and HiC loops49, snATAC-seq peak-to-peak co-accessible loops (r>0.45) within 10kbps of sSNP associated cCRE, ARL4C gene locus (chr2:234493041-234497081) has HiC linkages with sSNP region, gkmExplain importance scores for each base in 50-bp region surrounding sSNP for A1 and A2 alleles from Mic2 gkm-SVM model, the predicted TF binding site (FLI1) affected by the sSNP is shown at the bottom. C. Example plot for Mic2 sSNP (rs5995992). Top to bottom: snATAC-seq-derived pseudo-bulk tracks for each cell-type and Mic2 cluster (normalized by reads in TSS), Manhattan plot showing MDD lead SNP, SNPs in LD (r>0.8), and MDD sSNP (chr22: C>T; rs5995992) overlapping Mic2 candidate DAR (FDR=0.06) (peak-to-gene linkage: TOB2 gene), Mic2 candidate DARs and marker peaks in the plotted region, DLPFC neuronal and non-neuronal (NeuN+/-) H3K27ac and H3K4me3 ChIP-seq peaks merged across each subject25, DLPFC NeuN+/- promoter anchored HiC fragments and HiC loops49, snATAC-seq peak-to-peak co-accessibility (r>0.45) for cCRE within 10kbps of sSNP associated cCRE, TOB2 gene locus (chr22:41433494-41446801) with the most significant peak-to-gene linkage (r=0.8) with sSNP associated cCRE, No TF binding site was identified to be significantly disrupted by this sSNP.

### Molecular characterization of ExN1 and Mic2 clusters in depression

To relate molecular profiles of clusters with their likely functional roles, we compared promoter-accessibility with gene-expression from the Allen brain institute (ABI) snRNA-seq ^59^. ExN1 neurons had the highest similarity to layer 6 (L6) intratelencephalic (IT) Car3 neurons (AUC=0.98; Extended Fig 6A), which are transcriptionally similar to NR4A2-marked neurons ^60^ implicated in regulating stress-response ^61^. Likewise, compared to other excitatory neurons, ExN1 showed the highest TF motif accessibility and binding for NR4A2 (Fig 1H; Extended Fig 6B-C), also known as NURR1, which has been consistently linked to stress-response ^62–70^, and depression ^64,66,71,72^. Moreover, ExN1 marker TF, NR4A2 showed evidence for interactions with MDD-associated ExN1 TFs (Method: StringDB; direct interactions: bZIP: FOS, FOSB, bHLH: ASCL1; indirect: Zf: EGR1; Extended Fig 6D). Together, this presents an intriguing possibility for the role of coordinated TF-activities in NR4A2+ ExN1 neurons in regulating stress-related molecular alterations in depression.

On the other hand, between the two microglial clusters, Mic2 showed a significantly higher enrichment for GM than white matter (WM) microglial genes ^73^, (Extended Fig 6E-F). Likewise, Mic2 localized mostly to the GM when integrated with DLPFC spatial gene-expression dataset ^74^ (Extended Fig 6G). GM and WM microglia tend to be phenotypically different ^73^, and in a recent postmortem study on freshly-isolated microglia, only GM microglia showed differential gene-expression in MDD with downregulation of immune-related genes (referred to as, disease-associated microglia in depression (depDAM)) ^75^. Compared to Mic1, promoter-accessibility and gene-expression profiles of Mic2 reflected molecular status of depDAM (Extended Fig 6H-I), with significantly lower module scores for downregulated and higher for upregulated genes in MDD. While these patterns were not statistically different between groups, genes linked to downregulated Mic2 DARs in MDD significantly captured downregulated genes in depDAM (p-value=8.1e-05; one-sided Fisher’s exact test).

## Discussion

We examined chromatin accessibility with integrated gene-expression in ∼200,000 cells from the DLPFC. Our results indicate that nearly 75% of DARs identified across cell-type clusters were differentially closed in MDD individuals. These differences were found primarily in GM microglia (Mic2) and NR4A2+ deep-layer excitatory neuronal cluster (ExN1), which was also significantly enriched for the heritability of MDD GWAS SNPs.

In line with a previous cell-type-specific study examining PFC maturation ^76^, our study identified MDD-associated differentially accessible chromatin in deep-layer (L5/6) ExN1 neurons enriched for activity-dependent (bZIP) motifs. These neurons were also enriched for neurodevelopment-associated (bHLH/HTH) TF binding sites. Moreover, MDD-associated TFs from these families in ExN1 showed interactions with ExN1-specific marker TF, NR4A2. NR4A2 is an immediate-early factor ^62^ which responds to glucocorticoids ^67,77^, acute and chronic stress ^63–70^, and has been previously associated with depression ^64,66,71,72^. NR4A2 gene and protein levels were found to be decreased in bulk-tissue DLPFC from MDD individuals ^78^. Moreover, NR4A2 induction can have neuroprotective effects after exposure to neuropathological stress ^69^ and chronic inflammation-induced depression-like behaviors ^71^. The roles of NR4A2+ neurons in stress-response are consistent with observations in Nr4a2-null heterozygous mice showing significant reduction in cortical and limbic dopamine levels ^79^ and greater behavioral impairment associated with neuropsychiatric traits after exposure to developmental stress ^80^. This suggests that NR4A2+ ExN1 neurons may be more sensitive to stress-related perturbations in depression, possibly mediated by coordinated TF activities.

Furthermore, NR4A2+ ExN1 neurons were found to be most similar to L6 IT Car3 neurons ^71^, which share developmental and transcriptional similarities with NR4A2-marked Claustrum neurons ^60,81^. Chemogenetic silencing of these neurons attenuates stress-induced anxiety- and depression-like behaviors in rodents ^61^. Moreover, activity-dependent induction of NR4A2 can modulate AMPA-receptor mediated synaptic plasticity ^82^ whose alterations are strongly associated with depression ^83^.

In our study, NR4A2+ ExN1 neurons were also most significantly enriched for the heritability of MDD GWAS SNPs, especially in the marker cCREs, suggesting cell-intrinsic programs may be involved in regulating differential accessibility alterations. MDD-associated genetic variation in these neurons affected regulatory sites linked to genes involved in synaptic plasticity and communication, possibly impacting the sensitivity of ExN1 neurons to stress and depression.

In contrast to ExN1, a microglial cluster Mic2 in GM showed differentially closed chromatin associated with genes involved in phagocytosis- and immune-related pathways. Mic2 cluster also showed transcriptional similarities with a cortical GM microglia (depDAM), depicting down-regulation of genes involved in immune-response and phagocytosis, suggesting an overall immune-suppressed phenotype in MDD^75^. MDD-associated downregulation in immune-signaling pathways may be contributed by decreased binding activities of immune-regulatory TFs identified in this study, such as ETS domain and IRFs ^34,38^. Likewise, chronic stress-induced depression-like behavior in rodents has been associated with decreased interferon-response TF motif accessibility and transcriptional-repression in microglia ^84^. MDD-associated TFs identified in Mic2 were also among those depicting motif accessibility changes in response to disruption in microglia micro-environment and associated with neurodegenerative and psychiatric disorder genes ^28^. This suggests possible roles of surrounding tissue-environment ^28^ or microglia-neuron communication ^20,75^ in shaping regulatory landscape of microglia.

Previous work investigating tissue inflammation in depression produced contradictory findings, suggesting both increased ^85–87^ and decreased immune-activation ^10,75,88,89^, either peripherally or centrally. Consistent with our findings, bulk-tissue ^9^ and single-nucleus ^20^ RNA-seq in the DLPFC revealed downregulation of genes associated with inflammatory cytokines, microglia– activation, and immune-signaling pathways in MDD. Some studies in rodents ^84,90^ and humans ^91^ have likewise suggested suppression of microglial immune-response in chronic stress-induced disorders. Furthermore, recent reports provide evidence at gene-expression ^9,10,20,75^ and protein levels ^10,88,92^ for a non-inflammatory microglial phenotype in MDD.

It is possible that inflammatory changes taking place over time may eventually lead to widespread changes in chromatin accessibility of microglia, dysregulating TF binding and gene-expression and altering overall immune-homeostasis in MDD individuals. Additionally, MDD has been associated with alterations in microglial homeostatic functions ^10,88^, such as microglia-neuron communication ^20,75^. For example, early-life stress in mice induced disruption in microglia-associated pruning of excitatory synapses leading to aberrant adult stress-response ^93^. Whereas both microglia-suppressing and -stimulating properties associate with context-specific anti-depressant response ^94^. These results emphasize the need to further determine the influence of microglia-associated inflammatory as well as synapse-regulatory functions in context of depression.

Unlike excitatory neurons, marker cCREs specific to glial clusters were not significantly enriched for the heritability of MDD GWAS risk loci; however, examining variant-effects of individual MDD sSNPs pointed to disruption of specific gene-regulatory sites. Together, these results delineated regulatory mechanisms by which MDD-associated genetic and environmental factors may impact distinct cortical clusters (Extended Fig 7).

### Limitations

Sex-specific transcriptomic changes have been identified in depression ^20,95^, Here, we interrogated MDD-associated accessibility by accounting for sex of individuals but did not assess moderation effects of sex on phenotypic differences. Future studies hold promise to uncover chromatin regions possibly mediating sex differences in MDD.

The downstream interpretation of MDD-associated chromatin could be limited by the number of subjects or cells. Notably, the cohort-size of this study was comparable, if not larger, to other publicly-available postmortem single-cell brain studies ^18,24,76,96^. Due to cohort-size and overrepresentation of EUR population, we opted to use reference LD panels instead of in-sample LD. Consequently, our findings may not be applicable to other populations.

Finally, we adopted gkm-SVM models to predict allelic-impact of genetic variants on accessibility. Although the performance of our models is comparable to other studies ^18^, these models do not directly evaluate read-count differences between individuals ^97^.

## Conclusions

To our knowledge, our study is the first single-nucleus measurement of chromatin accessibility and integrated gene-expression in individuals affected by a psychiatric phenotype. Our study revealed MDD-associated accessibility alterations in the deep-layer excitatory neurons characterized by TF activity of NR4A2, which is known to regulate stress-response. NR4A2+ excitatory neurons were also most significantly enriched for the heritable risk for MDD, whereby the associated genetic variation disrupted regulatory sites linked to genes involved in synaptic communication. Additionally, a grey matter microglial cluster showed significant chromatin accessibility disruption in MDD, impacting binding of TFs known to mediate immune homeostasis. This rich resource will prompt future investigations into narrowing down the functional effects of MDD-associated regulatory sites and risk-variants using more targeted approaches, in attempts to further tease-apart the interplay of genetic and environmental factors contributing to MDD.

## Methods

### Post-mortem brain samples for snATAC-seq

We investigated a total of eighty-four human post-mortem dorsolateral prefrontal cortex samples obtained from the Douglas Bell Canada Brain Bank (www.douglasbrainbank.ca) and University of Miami Miller School of Medicine Brain Endowment Bank (https://med.miami.edu/programs/brain-endowment-bank). Clinical information was obtained using psychological autopsies, which were performed using proxy-based, structured and standardized interviews ^98^. This study was approved by the Douglas Hospital Research Ethics Board, and written informed consent from next of kin was obtained. The frozen histological grade samples of grey and white matter were dissected from the dlPFC (Brodmann Area 9) by expert neuroanatomists and stored at –80 °C. Cases met criteria for MDD and died by suicide during an episode of major depression, whereas controls were individuals who died of natural causes or in accidents and did not meet criteria for major axis I disorders. No differences were observed for the age of individuals (p-value=0.81, Case=44.6, Control=44.15). Although there was a difference between MDD and control subjects in the mean values for post-mortem interval (PMI; p-value=0.0037, Case=44.2, Control=33.4), no significant differences were observed for the pH values, which is a better correlate of tissue integrity (p-value=0.16, Case=6.5, Control=6.4; Supplementary Figure 2).

### Nuclei extraction, library multiplexing, and sequencing

We extracted nuclei using thick coronal sections (∼350um on average) as they were found to produce higher nuclei-to-debris ratio compared to thinner sections (data not shown). This also allowed for obtaining nuclei across cross-sections of GM and WM. Nuclei extraction from the homogenized brain tissue was performed as previously described ^99^ with some modifications (Supplementary note: Nuclei Extraction). We combined nuclei extracted from a male and female pair (either control or MDD) into one well of 10x microfluidic device (Supplementary note: Multiplexing and library construction) and sequenced the combined libraries together using Illumina NovaSeq 6000. These libraries were demultiplexed using 1000 Genomes based common variants and sex-specific chromatin accessibility (Supplementary note: Demultiplexing and assignment of sex). For each library, we obtained on an average 505M raw sequencing read-pairs with on an average 89.1% sequenced read pairs with mapping quality > 30 and having on an average 141.34 bp insert-size. Quality-metrics for each subject are provided in (Supplementary Figure 1, Supplementary Table 1).

### Doublet removal

10x fragment files were used as an input to ArchR [release_1.0.1]. High-quality cells were defined as those passing threshold criteria for the number of unique fragments per cell > 1000, TSSEnrichmentScore > 3.5 and fraction in mitochondrial reads < 10%. Cells detected as 10x gel-bead doublets or barcode multiplets originally identified in ^100^ and cells with possibly mixed genotype profiles detected using Vireo ^101^ were filtered out.

### Unsupervised iterative clustering

Genome-wide tile-matrix was generated by binning the genomic accessibility (GRCh38) into 500bp bins. Dimensionality reduction was then performed on the tile-matrix using the estimated iterative LSI procedure in ArchR and batch-effects were corrected with Harmony ^102^. Unsupervised graph-based clustering with a smart local moving algorithm was applied to addClusters(reducedDims=“Harmony”,dimsToUse=1:25,method=3,resolution=0.4,filterBias=TR UE). This first round of iterative clustering produced a total of 11 clusters, including 1 cluster of endothelial (C4), microglial (C5), astrocytic (C10), OPCs (C11), inhibitory neurons (C6), 3 oligodendrocyte clusters (C1-3), and 3 excitatory neuronal clusters distinguished by upper (Cluster 7), middle (C8), and deep cortical layers (C9), which were then combined for subsequent clustering within broad cell-types. For each broad cell-type, we re-computed LSI (15,000 varFeatures and 1:20 dimsToUse) and retained clusters with at least 50 cells. The second round of clustering was performed at multiple resolutions (0.2,0.4,0.6), retaining the clustering resolution that produced the highest median of mean Silhouette scores across clusters. For larger cell-types (oligodendrocytes and excitatory neuronal cell-types) for which Silhouette scores could not be computed (>70,000 cells), we used clustering resolution of 0.6.

### Quality-assessment of clusters

The quality of identified clusters produced after the first and second rounds of clustering was assessed based on: a) uniform percentage contribution from subjects, batches, sexes, and conditions in each cluster, b) having at least 10 marker genes based on ArchR GeneScore matrix (Wilcoxon test, FDR < 0.05 & Log2FC > 0.5). This retained 11 out of 15 clusters after the first round and 38 out of 40 clusters after second round of clustering for downstream analysis.

### Identification of chromatin cis-regulatory elements (cCREs)

To identify reproducible sets of cCREs capturing biological variability across cell-types and clusters, cells belonging to same cell-type and cluster were first partitioned into four non-overlapping pseudobulk aggregates whereby each replicate had uniform contributions from subjects belonging to each sex and condition. Open chromatin regions (OCRs) were called using MACS2 ^103^ on each pseudobulk replicate available for each cell-type and cluster using addReproduciblePeakSet(excludeChr=“chrM”, maxPeaks=300000, cutOff=0.1) in ArchR. Finally, OCRs reproducible in at least two of the four replicates in each broad cell-type and cluster were retained. Next, we generated a non-overlapping and fixed-with peak-set (501bp) separately at cell-type and cluster levels using iterative overlap peak merging and removal (IPR) procedure in ArchR. This retains the most significant OCR (based on MACS2 significance) among those overlapping across clusters. OCRs accessible in at least 1% of cells resulted in approximately 600 and 800 thousand non-overlapping cCREs at broad cell-type and cluster resolution, respectively. Majority of broad cell-type cCREs (86%) overlapped with cluster-resolved cCREs. Conversely, a much smaller percentage of cluster cCREs (64%) overlapped with broad level cCREs. Notably, most cluster cCREs (95%) were located within 5000bp neighborhood of broad cCRE coordinates. Finally, we computed per-cell Tn5 insertion counts in each cell with respect to the above-mentioned non-overlapping cCREs from broad cell-type and cluster resolution, which were then used for downstream analyses.

### Marker cCREs

Cell-type- and cluster-specific marker cCREs were identified using the respective Tn5 insertion count matrices. For this, getMarkerFeatures() in ArchR was used with Wilcoxon test (FDR < 0.05 & Log2FC > 0.5). Majority of broad cell-type-specific marker cCREs (77%) overlapped with cluster-specific marker cCREs. Conversely, a much smaller number of cluster marker cCREs (36%) overlapped with the broad marker cCREs. About 67% of cluster marker cCREs were within 5000bp of broad marker cCRE coordinates. Three (ExN7, InN3, and Oli6) of the 38 clusters were potentially low-quality as they had very few marker cCREs (<5) compared to all others.

### Cluster annotation

Clusters were annotated based using several approaches: a) pseudo-bulk accessibility profiles (normalized by reads in TSS) at cell-type marker genes plotted using ArchR plotBrowserTrack function, b) assessing gene regulatory activity of cell-type marker genes in BrainInABlender ^104^ (Species==“Human”) for every cluster using ArchR addModuleScores() function, and c) measuring gene activity profiles with ArchR GeneScore matrix (Wilcoxon test; FDR < 0.05 & Log2FC > 0.5) and manually assessing the enrichment of following marker genes:

Neurons: *SNAP25, RBFOX3*; Excitatory neurons: *SATB2, SLC17A7, SLC17A6*; Excitatory neuronal cortical layers (upper to lower): *CUX2, RORB, PCP4, BCL11B, FEZF2*; Inhibitory neurons: *GAD1, GAD2, SLC32A1*; Inhibitory neuronal subtypes: *LHX6, PVALB, SST, ADARB2, VIP, LAMP5* Macrophage/microglia: *MRC1, CSF1R, CX3CR1*; Endothelial: *CLDN5, SRGN, FLT1*; Astrocytes: *AQP4, SLC1A2, GFAP*; OPCs: *HAS2, CSPG4, OLIG1*; OPCs: *HAS2, CSPG4*, and Oligodendrocytes: *MBP, OPALIN, PLP1, MAG, MOG*.

### Validation of cell type clusters

#### Enrichment of Fluorescence-activated nuclei (FAN)-sorted cell-type marker peaks

We thoroughly validated snATAC-seq cell-type and cluster profiles using various strategies. First, we examined enrichment overlap of marker cCREs from FAN-sorted bulk ATAC-seq cell-types ^105^ in cluster-specific marker cCREs defined by our snATAC-seq data using ArchR peakAnnoEnrichment(cutOff =“FDR < 0.05 & Log2FC > 0.5”) and plotted FANS-sorted cell-type cCREs with the highest enrichment for each cluster using plotEnrichHeatmap(enrichRegions, n = 1).

#### Comparison with external snATAC-seq dataset

Second, using Signac ^106^, cell-types identified in our data were compared to the cell-types available in PFC snATAC-seq ^24^. Specifically, for each cell in our cohort, fragment counts were computed using peaks identified in the external PFC snATAC-seq dataset. Next, cell-type labels from the external snATAC-seq were transferred onto our data using cell-by-peak fragment count matrix and using FindTransferAnchors(reduction=“lsiproject”, dims=2:20) function in Seurat ^109^.

#### MetaNeighbor

Finally, we assessed the concordance between cell-type and clusters in this dataset with multiple snRNA-seq datasets. For this, we used the top 3000 most variable features to create gene-expression classifiers for each of the cluster defined in these snRNA-seq datasets ^20^ ^59^ ^107^. Using MetaNeighbor ^108^, we predicted their similarities (AUCROC plots; Extended Figure 2C-D) to snATAC-seq clusters based on promoter-accessibility (<2kbp to gene TSS) computed using GeneActivity() function in Signac ^106^

### Integration with snRNA-seq dataset

Prior to integration, cells with uncertain (“Mixed”) annotation in MDD snRNA-seq data ^20^ were removed. First, an unconstrained integration was performed using snRNA-seq gene-expression as reference and snATAC-seq GeneScore matrix as query to derive prediction scores and assess concordance between cell-types. As we observed high similarity between cell-type labels between two datasets (Extended Fig 1A-B), we individually applied constrained integration within broad cell-types across two datasets and subsequently imputed gene expression from snRNA-seq to snATAC-seq cells using addGeneIntegrationMatrix(reducedDims=“Harmony”, addToArrow = TRUE, dimsToUse=1:20) function in ArchR.

### Integration with spatial gene-expression dataset

snATAC-seq microglia clusters (Mic1 and Mic2) were mapped onto 10x Visium spatial gene-expression dataset comprising of male and female DLPFC tissue sections ^74^ using Seurat ^109^. Briefly, SCTransform() was used on promoter-accessibility and integrated snRNA-seq gene-expression for snATAC-seq cells mapping to microglial clusters. Microglia cluster labels were then transferred using the top 30 principal components (PC) with FindTransferAnchors() function and predicted spatial positions were visualized using SpatialFeaturePlot() in Seurat.

### Differential accessibility analysis

Differential accessibility analysis was performed in every broad cell-type and cluster using limma-voom ^110^ implemented in muscat ^111^. First, pseudo-bulk accessibility profiles were generated by aggregating counts across cells for each cCRE for each subject within the broad cell-types and clusters. One female was omitted prior to differential analysis due to discordance between the sex identified and reported (Supplementary note). Batch, sex, age, PMI, and the total number of cells contributed by each subject were added as covariates in the limma-voom differential accessibility model. A series of quality-control measures for differential accessibility analyses were taken: a) Filtering out subjects with very low cell contribution (less than 10 cells in broad cell-types or 5 cells in clusters), b) Filtering out potential outlier subjects based on peak-count matrix using isOutlier(type = “lower”, nmads = 3) function in muscat, c) Filtering out cCREs with consistently low counts across multiple subjects using filterByExpression() function from edgeR ^112^. Finally, for each broad cell-type and cluster, we retained cCREs accessible in more than 1% and 3% of cells, respectively, on an average in either cases or controls. Differentially accessible regions (DARs) with abs(logFC) > log2(1.1) and FDR-adjusted local p-value < 0.05 was used for all the analysis, except GWAS based analysis for which we defined candidate DARs (abs(logFC) > log2(1.1) and FDR < 0.2).

### Differential motif analysis

DARs were first partitioned by the direction of their association with MDD and were then provided as inputs to Homer. GC-content matched background regions were automatically selected by Homer and motifs were scanned within 200bp region of the peak center using findMotifsGenome.pl function with default parameters. We only assessed the “known” motifs for downstream results. Additionally, ChromVar ^39^ based GC content and mean accessibility biases-corrected motif accessibility deviations z-scores were computed at single-cell level with Cisbp motif database using addDeviationsMatrix() function in ArchR. Maker TFs for each cell-type and cluster were defined as those with significantly higher motif accessibility in a cluster compared to all others using Wilcoxon test (FDR < 0.05 & MeanDiff > 1.5) with getMarkerFeatures() default function in ArchR. MDD-associated TFs were also identified using getMarkerFeatures(useGroups = “Case”, bgdGroups = “Control”) after adding PMI, age, and total number of fragments as bias variables.

### TF motif footprinting

We generated TF binding profiles by aggregating Tn5 insertion counts across all the motif matching peaks using Cisbp motif database and subtracting Tn5 insertion sequence bias before plotting. Specifically, for cell-type and cluster-specific marker TFs, we generated a maximum of 5 pseudo-bulk aggregates (default parameter) representing each cluster. For MDD-associated TFs, we created 10 pseudo-bulk replicate profiles from each group using addGroupCoverages(groupBy=”Condition”, maxReplicate=10) in ArchR.

### Gene regulatory networks

We prioritized MDD-associated TFs for constructing gene regulatory networks (Supplementary Figure 5E). Briefly, among the topmost significantly enriched TF motifs (FDR<0.05) in cluster-specific DARs, we selected the top two TFs with the highest median gene-expression using snRNA-seq integrated expression. Between these two TFs, we prioritized the one with the most significant difference (FDR<0.05) between case and control cells using ChromVAR. For ExN1 gene-regulatory network, following DARs (FDR<0.05, abs(logFC) >log2(1.1)) were used, but for Mic2 network, we selected topmost significant DARs (FDR<0.01, abs(logFC) >log2(1.1)). To improve readability of graphs, we only plotted DARs mapping to ArchR-defined non-coding elements, including promoters, introns and distal regions and excluded exons. The edges in these networks are directed from DAR nodes to gene nodes and are drawn whenever the corresponding DAR contained a TF binding site and: (a) connects any DAR (promoter, introns, or distal) to corresponding target-genes through peak-to-gene linkages (r>0.45); and (b) connects promoter DARs to their nearest gene. Directed gene-regulatory networks were visualized using the Fruchterman-Reingold layout algorithm with the R package igraph (v 1.2.6). The nodes in green represent DARs enriched for individual TF binding sites directed towards putative target-genes colored according to MDD associated gene or MDD DEG status. For MDD-associated genes, we compiled list of potential risk-genes in MDD GWAS ^11–13,72,113,114^, Hi-C based analysis (H-MAGMA; FDR<0.1) ^115^ and PsyGeNET depressive disorders genes ^26^. For MDD DEGs, we used bulk (FDR<0.1) ^116^ and snRNA-seq DEGs ^20^ obtained from the meta-analysis of both sexes (FDR<0.1, abs(log2FC) > log2(1.1), signs > 0).

### Functional interpretation of differential results

- **Co-accessible peaks and peak-to-gene linkages** Co-accessibility of all pairs of cCREs at cell-type and cluster levels were computed using via addCoAccessibility(maxDist=5e+05) function in ArchR. Moreover, using MDD snRNA-seq integrated expression, for all pairs of cCREs and genes, we computed the correlations between cCRE-accessibility and gene-expression, using addPeak2GeneLinks(maxDist = 5e+05) in ArchR. We pruned the resulting peak-to-peak co-accessibility and peak-to-gene linkage matrices to retain all significantly correlated pairs (r>0.45), as defined in ArchR. Additionally, we retained genes that were most significantly correlated (lowest FDR values) with cCREs associated with MDD sSNPs using peak-to-gene linkages (Supplementary Table 12).
- **Gene ontology (GO) pathways** Gene ontology pathways were identified for DAR-linked genes (r>0.45) in each cluster using enrichGO(minGSSize=5, pAdjustMethod=“BH”, pvalueCutoff=0.05, ont=“ALL”) with clusterProfiler ^117^. We also split genes according to differentially more or less accessible DARs and performed pathway enrichment analysis using more relaxed parameters: enrichGO(pAdjustMethod=“BH”, pvalueCutoff=0.2).
- PsyGeNET For examining the association of DAR-linked genes with psychiatric phenotypes, we used psygenet2r package with database=“ALL” and default parameters associated with PsyGeNET database ^26^. The total number of gene-disease associations and evidence index for each psychiatric disorder was computed using geneAttrPlot() with default parameters.
- **StringDB** Protein-protein (PP) interactions were extracted from “full String network” of organism= “Homo Sapiens.” The minimum required interaction score was set to “high” (i.e., 0.7) and the maximum number of interactions allowed in first and second shells of the network were limited to 20 and 5 interactions, respectively. The fully-connected PP interaction network was generated by removing the disconnected nodes.
- **cCRE overlap analysis** To compute overlap of PsychEncode H3K27ac and H3K4me3 histone modification peaks consolidated from cortical cells ^25^ with marker cCREs, cCREs in peak-to-gene linkages (r>0.45), and MDD DARs, we used subsetByOverlaps() in IRanges R package with default parameters.
- **Gene overlap analysis** To assess enrichment of snRNA-seq DEGs, GWAS risk-genes, PsyGeNET depressive disorder genes in DAR-linked genes at broad and cluster levels, we used one-sided Fisher’s exact test in newGOM() GeneOverlap R package [v.1.36.0; http://shenlab-sinai.github.io/shenlab-sinai/] and plotted FDR-corrected p-values using drawHeatmap(go.obj, adj.p = T,log.scale = T, ncolused = 9).
- **Module scores** Gene-expression module scores (average weighted expression of genes at single-cell level) were calculated using Seurat AddModuleScore() function ^109^ for DAR-linked genes and TF target-genes. These scores were compared between MDD versus control cells (Wilcoxon test) using the most similar snRNA-seq defined clusters ^20^ to ExN1 and Mic2 snATAC-seq clusters. Both ExN16_L56 and ExN20_L56 snRNA-seq clusters showed correspondence to ExN1 (Extended Figure 2C); however, similarly to ExN1, ExN16_L56 also showed the highest similarity ^20^ with ExN L6 IT Car3 neurons in the ABI snRNA-seq ^59^, and had the highest mean expression for *NR4A2* (Extended Figure 2E), a cell-defining TF marker for ExN1. On the other hand, snRNA-seq data defined only one microglial cluster, which was used for the assessment of Mic2 gene-expression module scores. In addition, we examined module scores using promoter-accessibility and snRNA-seq integrated gene-expression for molecular characterization of microglia. For GM versus WM comparison, we used top 25 differential genes between GM and WM microglia (sorted by fold change) ^73^. For comparison with MDD-associated microglia (depDAM), we used all the genes that were upregulated or downregulated in MDD ^75^.

### Heritability enrichment

To calculate heritability enrichment of GWAS SNPs within cluster-specific marker cCREs and MDD-associated chromatin, we applied stratified LDSC regression model (v 1.0.1) to GWAS summary statistics for MDD ^11^ ^13^ and other complex traits ^118^ ^119^ ^120^ ^121^ ^122^ ^123^ ^124^. Cluster-specific marker cCREs were retained at FDR<0.05 & Log2FC>0.5. For MDD-associated chromatin, we used candidate DARs (abs(logFC) > log2(1.1) & FDR < 0.2) to obtain topmost dysregulated cCREs representing most of the clusters. These cCREs were converted from hg38 to hg19 coordinates using the UCSC liftover tool (v377). Cell-type specific partitioned heritability analysis was computed using a full baseline-LD model and after adding LD scores estimated on matching cCRE set combined across all clusters as background. Finally, one-sided p-values were computed based on the enrichment z-scores for each annotation relative to the background for every trait.

### Heritability Estimator from Summary Statistics (HESS)

We used HESS ^43^ to estimate local heritability for MDD-associated variants ^11^ in LD blocks defined on 1000 Genomes ^125^. This identified LD blocks significantly enriched (p<0.05) for MDD heritability. We then tested the enrichment of cluster-specific candidate DARs within MDD-associated heritable LD blocks identified above. For this, we compared cluster-specific overlap of candidate DARs in significantly heritable LD blocks for MDD (n=90) with per-cluster null distribution of the overlap of chromosome-matched cCREs within any LD block permutated 10,000 times to derive an empirical p-value of significance (p<0.05). Finally, both ExN1-specific marker cCREs and candidate DARs whose peak-coordinates were located specifically within significant MDD heritable LD blocks were input to Homer motif analysis (as described above). TF families enriched (q<0.05) commonly in both ExN1 marker cCREs and candidate DARs were identified. The occurrence frequency of enriched TF families was plotted using worlcloud2(v. 0.2.1) based on the number of individual TF members (belonging to each family) that were significantly enriched in ExN1-specific cCREs.

### Enrichment of DLPFC expression and splicing QTLs in cCREs

To perform e/sQTL enrichment in cCREs, we used binomial test to assess whether uniformly processed DLPFC eQTL and sQTLs (FDR<0.05; Supplementary method) are significantly enriched within cCREs. First, eQTL and sQTL SNPs located within 501-bp cCREs were identified. Next, the expected number of eQTL and sQTLs in cCREs were computed as the product of (1) cCRE coverage with respect to hg38 genome; and (2) total number of QTL SNPs at FDR < 0.05 (∼3,000,000 eQTL and ∼2,450,000 sQTL SNPs). Furthermore, binomial enrichment test p-values are calculated for each QTL type and cCREs. Here, success rate, number of trials, and number of successes for the binomial test were set to the cCRE coverage percentage, total number QTL SNPs, and the number of QTL SNPs observed in cCREs, respectively.

### Functional impact of MDD-associated genetic variation in each cluster

To identify allelic-effects of MDD-associated GWAS SNPs and those associated by LD (r>0.8) and fine-mapping (Supplementary methods) on chromatin accessibility of each cluster, we adapted three complementary gapped kmer support vector machine (gkmSVM) models ^46^ ^47^ ^48^. Briefly, for each of the snATAC cluster, gkmSVM models were trained on top 60,000 OCRs (whenever available) identified in each cluster by MACS2 (ranked by FDR) to predict whether 1001bp reference genome sequences were accessible (i.e., cCRE) or inaccessible (i.e., GC-content and chromosome-matched sequences) for the corresponding cluster. We specifically focused on MDD SNPs located in either marker cCREs or candidate DARs of each cluster (e.g., ExN1) and its respective cell-type (e.g., ExN) to assess the effects of MDD sSNPs at cluster-refined resolution. Using cluster-specific trained models, we then identified significant MDD SNPs (sSNPs) altering chromatin accessibility of 201bp neighborhood of a cCRE. Moreover, we partitioned sSNPs identified per cluster to high, medium and low confidence sets reflecting significance of accessibility disruption in its immediate local neighborhood (<201bp). Putatively disrupted TF binding sites were determined by matching affected allele sequence with Cisbp motif database in TOMTOM ^126^.

### Multi-modal Visualization

For interpreting functional targets of MDD sSNPs, we used snRNA-seq integrated peak-to-gene linkages (r>0.45) and snATAC-seq based peak-to-peak co-accessibility (r>0.45) using all the co-accessible loops within 10kbp region of cCREs associated with MDD sSNPs. We further overlayed PsychEncode DLPFC histone modification peaks (H3K27ac, H3K4me3) merged across subjects ^25^ and DLPFC NeuN+/- promoter-interacting Hi-C linkages (10kbp resolution) ^49^ (Supplementary note: PsychEncode data).

---

**Extended Figure 1.**
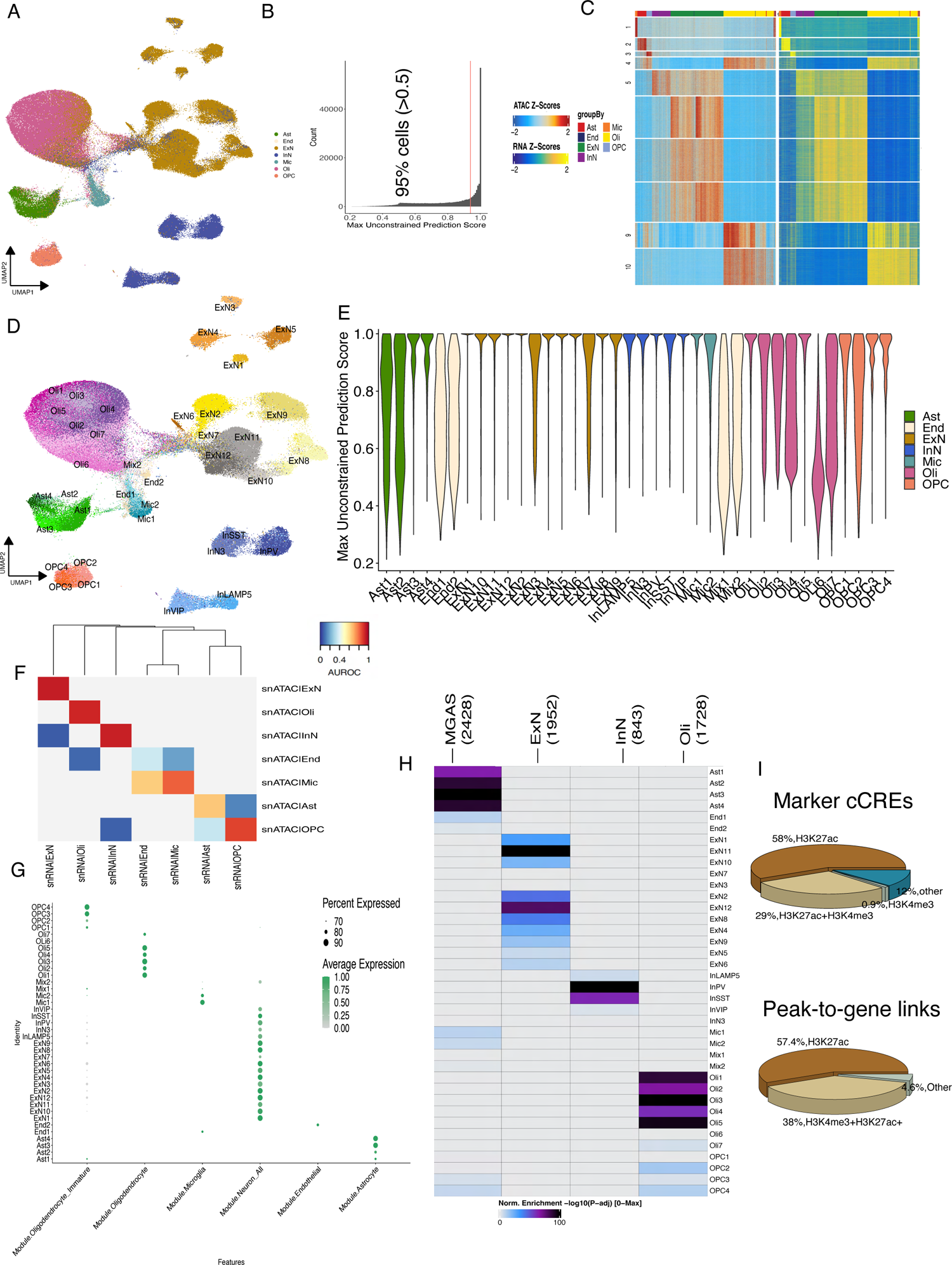
Annotation of snATAC-seq cell-types and clusters. A. UMAP plot based on chromatin accessibility in cCREs identified at cluster resolution and colored by cell-type labels predicted from MDD snRNA-seq from the same samples. B. Histogram shows MDD snRNA-seq cell-type label-transfer prediction scores across snATAC-seq cells (median prediction score: 0.95; in red). C. K-nearest neighbors (KNN; k=10) based on snATAC-seq peak-accessibility z-scores (left) and snRNA-seq integrated gene-expression z-scores (right) for significant peak-to-gene linkages (r>0.45) grouped by snATAC-seq cell-types. D. UMAP plot based on chromatin accessibility in cCREs identified at cluster resolution and colored by snATAC-seq clusters. E. Distribution of MDD snRNA-seq cell-type label-transfer prediction scores across snATAC-seq cells in each cluster. F. MetaNeighbor best hit plot depicting correspondence (AUROC) between cell-types in MDD snATAC-seq and snRNA-seq datasets. G. Dotplot depicting module scores (size of dots represents percentage of cells) in each snATAC-seq cluster computed based on gene-activity (ArchR GeneScore matrix) for cell-type marker genes catalogued in BrainInABlender104. H. Heatmap shows the overlap enrichment of the most significant fluorescence-activated nuclei (FAN)-sorted bulk ATAC-seq cell-type marker peaks in snATAC-seq cluster-specific marker peaks (Hypergeometric test; FDR<0.05 & Log2FC > 0.5; n=1; MGAS: Microglia and Astrocytes; ExN: Excitatory neurons; InN: Inhibitory neurons; Oli: Oligodendrocyte-lineage cells). I. Overlap of PsychEncode ChIP-seq histone modification peaks (H3K27ac, marks active enhancers; H3K27ac+H3K4me3: marks promoters) from cortical cells with cluster-specific marker cCREs (top) and cCREs in significant peak-to-gene linkages (r>0.45) (bottom).

**Extended Figure 2:**
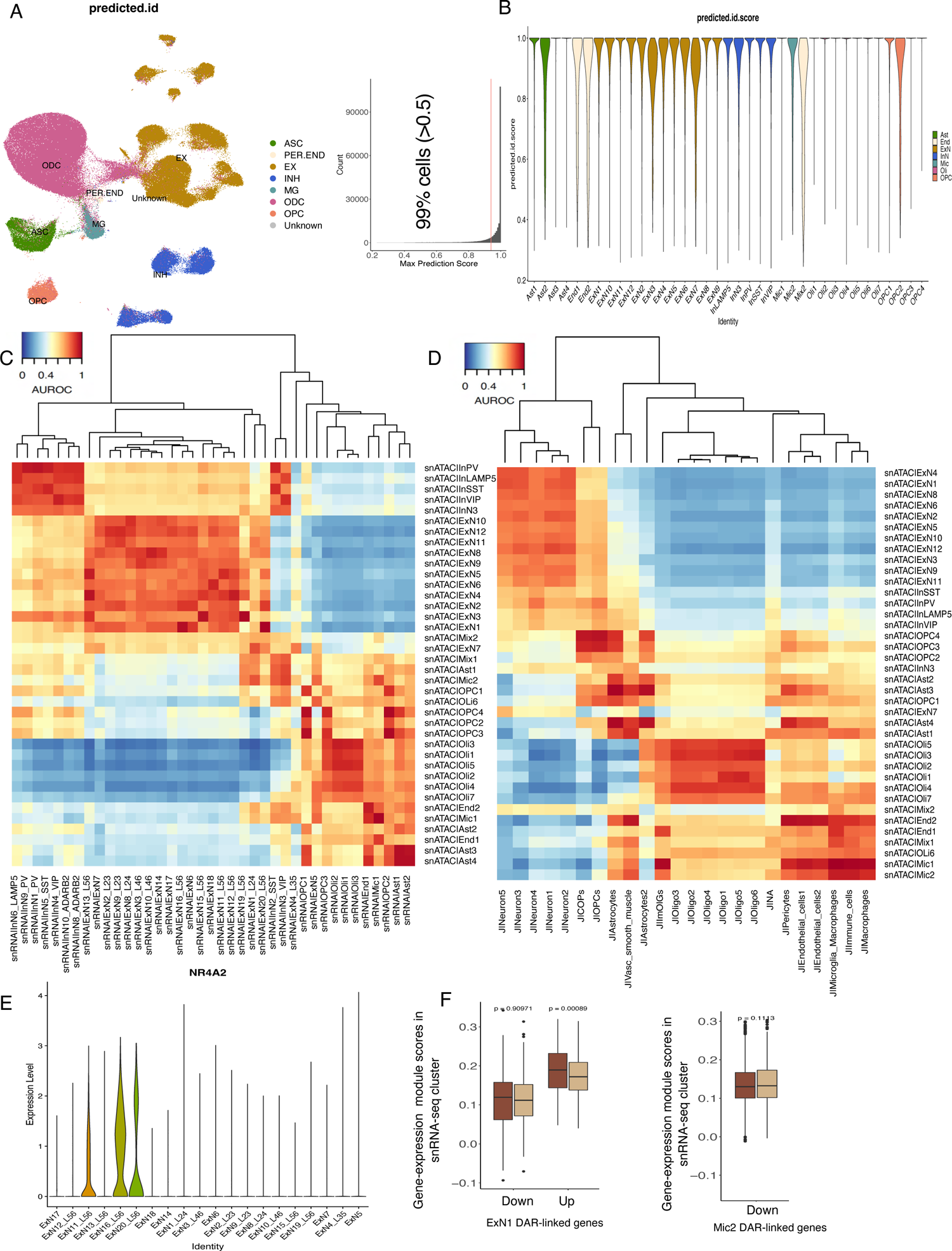
Comparison of snATAC-seq with published single-cell datasets. A. UMAP plot based on chromatin accessibility in cluster resolution cCREs colored by cell-type labels predicted from the reference PFC snATAC-seq24. Histogram shows reference label-transfer prediction scores across cells in our data (median prediction score: 0.99; in red). B. Distribution of reference cell-type label-transfer prediction scores across snATAC-seq cells in each cluster. C-D. MetaNeighbor AUROC plots show correspondence of MDD snATAC-seq gene promoter-accessibility with C. MDD snRNA-seq clusters20 and D. White matter postmorterm human brain tissue snRNA-seq clusters107. E. Violin plot shows NR4A2 gene expression in excitatory neuronal clusters identified in MDD snRNA-seq20. F. Gene expression module score differences in MDD versus controls in corresponding snRNA-seq clusters. NR4A2+ excitatory neuronal cluster (ExN16_L56) for ExN1 DAR-linked genes (left) linked to significantly less accessible (down; FDR<0.05) or more accessible DARs in MDD (up; FDR<0.05) and gene expression module score differences in snRNA-seq Mic cluster for genes liked to differentially less accessible Mic2 DARs (FDR<5%; right).

**Extended Figure 3:**
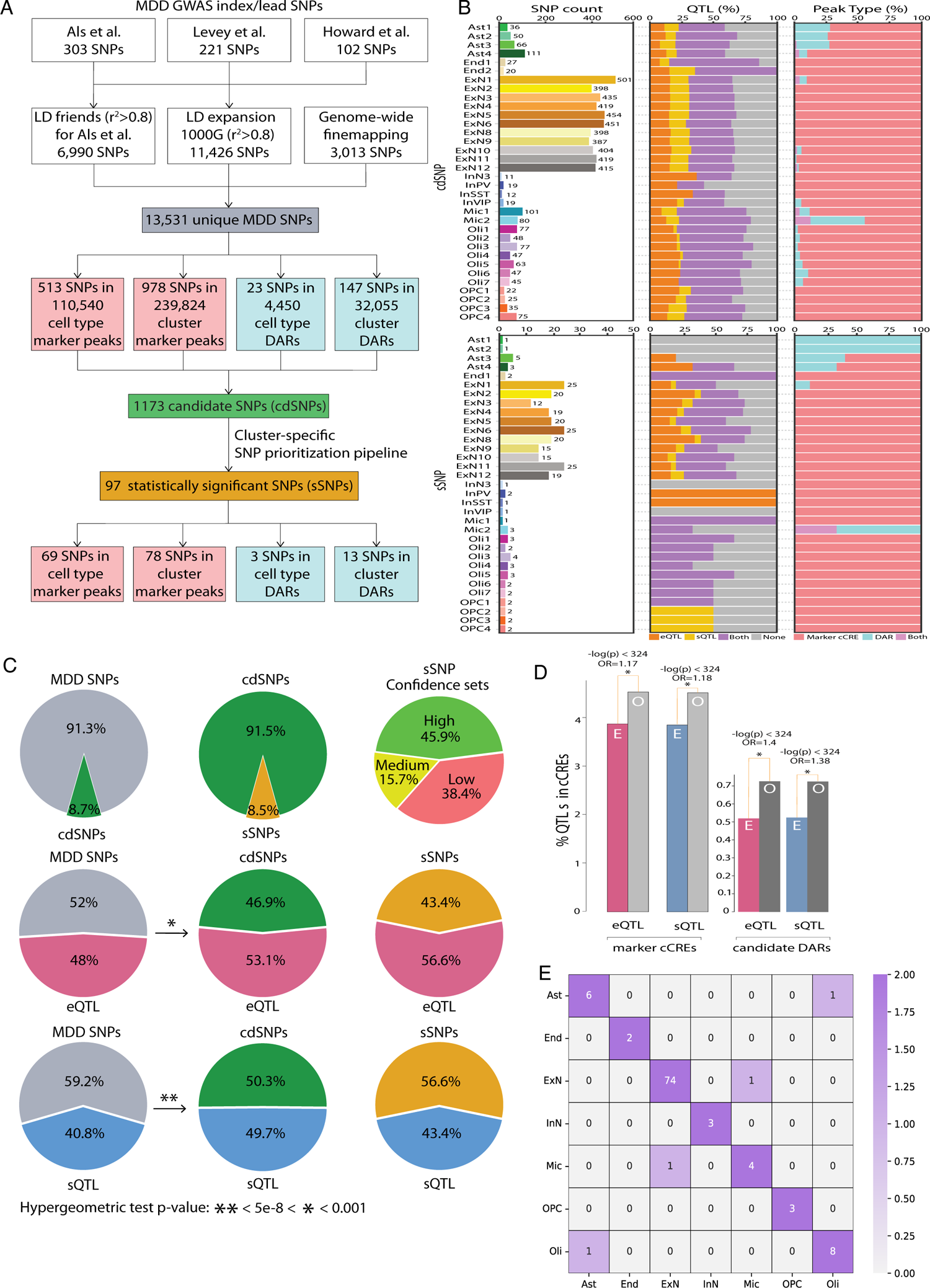
MDD risk variants identified in each cluster. A. Flowchart for identification of significant MDD SNPs (sSNPs) with allelic-effects on cluster-specific chromatin accessibility predicted to be significant across all three cluster-specific gkmSVM models (ISM, deltaSVM, gkmexplain) B. Stacked bar plots show overview of cdSNPs (top) and sSNPs (bottom) in each cluster (left to right: counts in each cluster, % SNPs that are significant eQTLs and sQTLs in the DLPFC (FDR<0.05), % SNPs mapping to cluster-specific marker cCREs and candidate DARs. C. Pie charts depicting (first row) percentage of cdSNPs in MDD-associated SNPs, percentage of sSNPs in cdSNPs, and sSNPs categorized based on accessibility disruption confidence sets; (second row) percentage of significant DLPFC eQTLs in MDD-associated SNPs, cdSNPs, sSNPs. Arrow represents significant enrichment (p<0.001; hypergeometric test) of eQTLs in cdSNPs compared to all MDD-associated SNPs; (third row) depicting percentage of sQTLs in MDD-associated sSNPs, cdSNPs, sSNPs. Arrows represents the significant enrichment (p<5e-08; hypergeometric test) of sQTLs in cdSNPs compared to all MDD-associated SNPs. D. DLPFC eQTLs and sQTLs were significantly enriched (binomial test, p<0.05) in marker cCREs and candidate DARs compared to the expected genomic coverage. E. Heatmap showing sSNP overlap across cell-types. Each cell in the plot is scaled with respect to the total number of sSNPs available for the cell type in the column.

**Extended Figure 4:**
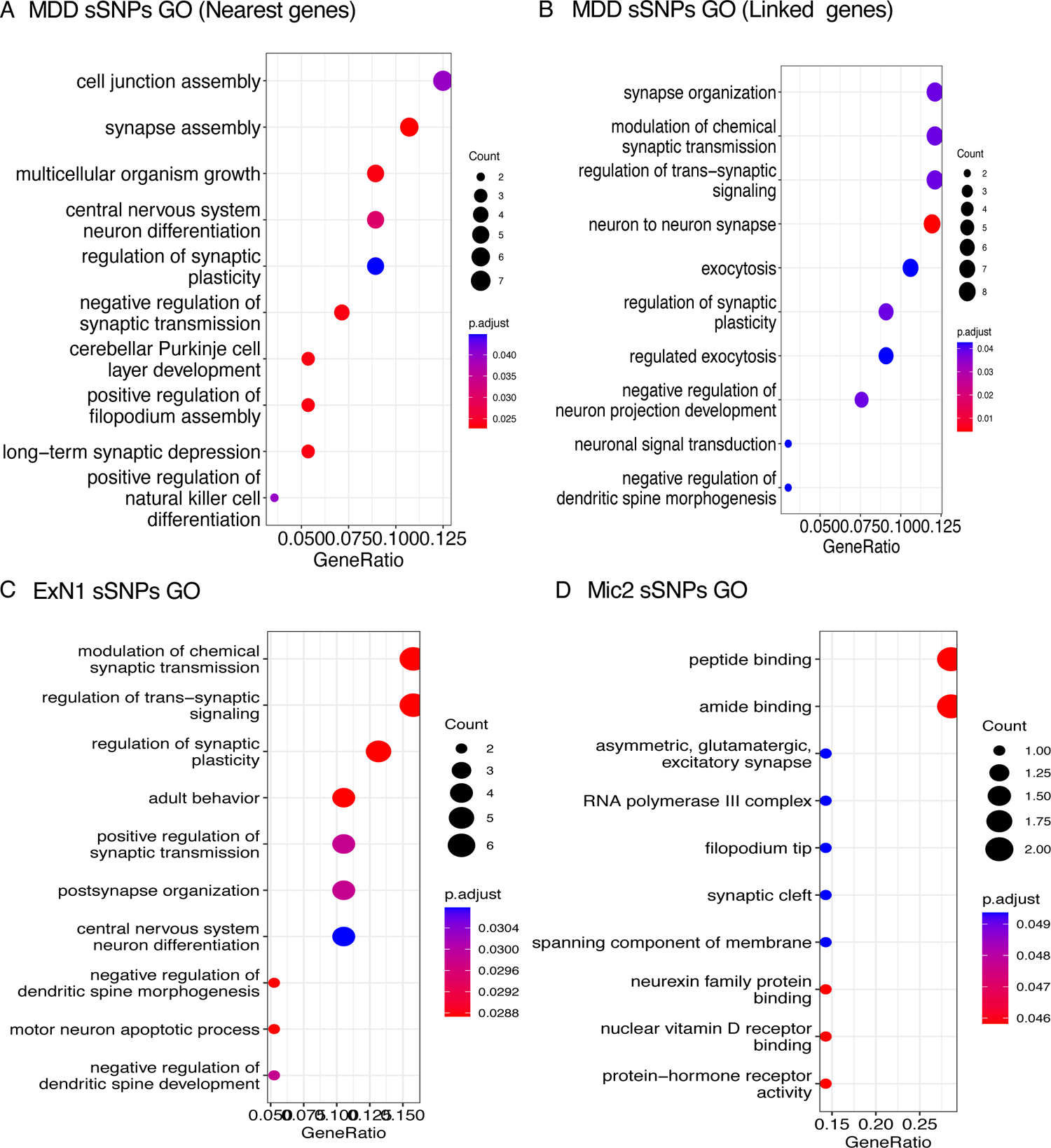
Gene ontology (GO) pathways associated with MDD risk variants. GO pathways enriched (FDR<0.05) in genes associated with MDD sSNPs A. nearest genes, B linked genes (peak-to-gene linkages, r>0.45). C. GO pathway enrichment (FDR<0.05) for ExN1 sSNP associated genes (nearest, linked, and eQTLs). D. GO pathway enrichment (FDR<0.05) for Mic2 sSNP associated genes (nearest, linked, and eQTLs).

**Extended Figure 5.**
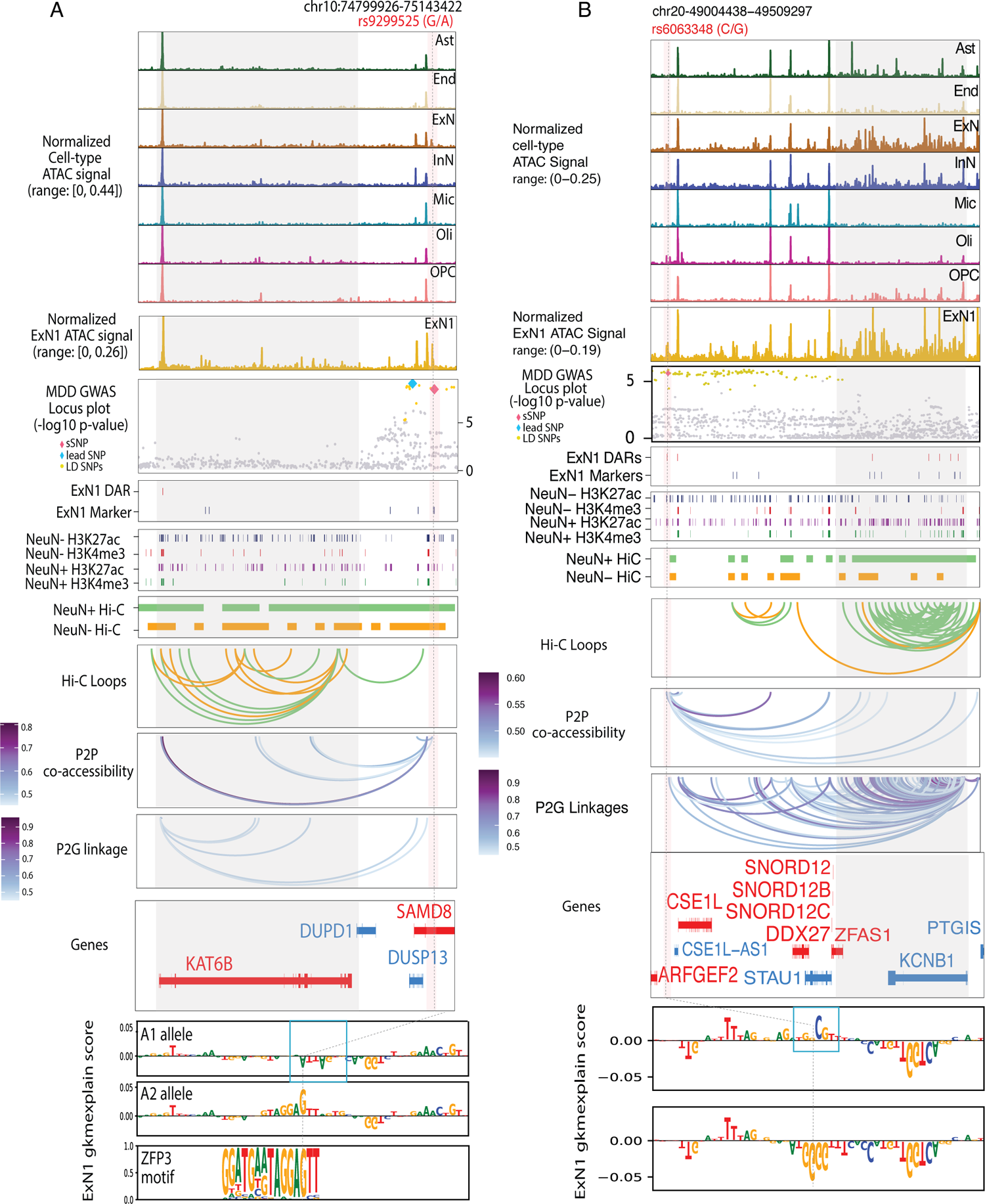
Examples of MDD sSNPs with allelic-effects exclusively on chromatin accessibility in ExN1 cluster. A. Example plot for ExN1 sSNP (rs9299525). Top to bottom: snATAC-seq-derived pseudo-bulk tracks for each cell-type and ExN1 cluster (normalized by reads in TSS), Manhattan plot showing MDD sSNP (chr10: G>A; rs9299525) overlapping ExN marker cCRE (peak-to-gene linkage: KAT6B gene), ExN1 candidate DARs and marker cCREs within the plotted region, DLPFC neuronal and non-neuronal (NeuN+/-) H3K27ac and H3K4me3 ChIP-seq peaks merged across each subject25, DLPFC neuronal and non-neuronal (NeuN+/-) H3K27ac and H3K4me3 ChIP-seq peaks merged across each subject25, DLPFC NeuN+/- promoter anchored HiC fragments and HiC loops49, snATAC-seq peak-to-peak co-accessible loops (r>0.45) within 10kbps of sSNP associated cCRE, KAT6B gene locus (chr10:74824927-75032624) with the most significant peak-to-gene linkage (r=0.4) with cCRE associated with sSNP, gkmExplain importance scores for each base in 50-bp region surrounding sSNP for A1 and A2 alleles from ExN1 gkm-SVM model corresponding, the predicted TF binding site (ZFP3) affected by the sSNP is shown at the bottom. B. Example plot for ExN1 sSNP (rs6063348). Top to bottom: snATAC-seq-derived pseudo-bulk tracks for each cell-type and ExN1 cluster (normalized by reads in TSS), Manhattan plot showing MDD sSNP (chr20: C>G; rs6063348) overlapping ExN1 candidate DAR (FDR=0.15) (peak-to-gene linkage: KCNB1 gene), ExN1 candidate DARs and marker cCREs within the plotted region, DLPFC neuronal and non-neuronal (NeuN+/-) H3K27ac and H3K4me3 ChIP-seq peaks merged across each subject25, DLPFC NeuN+/- promoter anchored HiC fragments and HiC loops49, snATAC-seq peak-to-peak co-accessible loops (r>0.45) within 10kbps of sSNP associated cCRE, KCNB1 gene locus (chr20-49293394-49484297) with the most significant peak-to-gene linkage (r=0.4) with cCRE associated with sSNP, No TF binding site was identified to be significantly disrupted by this sSNP.

**Extended Figure 6.**
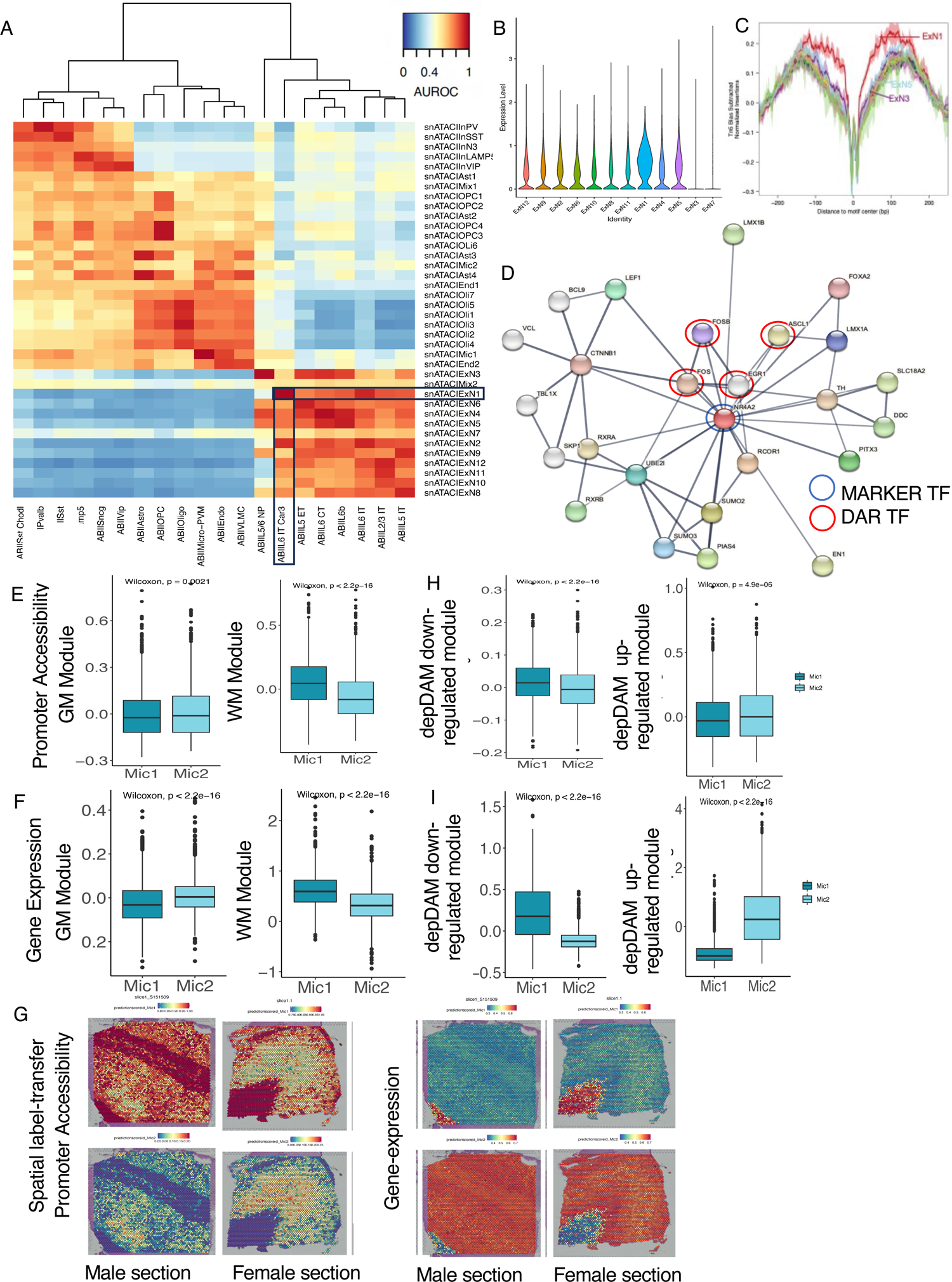
Functional characterization of ExN1 and Mic2 clusters. A. MetaNeighbor AUROC plot shows the strongest correspondence of snATAC-seq ExN1 cluster with ExN layer 6 IT Car3 neurons in Allen brain institute (ABI) snRNA-seq from the motor cortex (M1)59. B. Violin plot depicts promoter-accessibility for NR4A2 gene in ExN clusters. C. Tn5 bias subtracted NR4A2 motif footprints aggregated in each of the ExN clusters. D. StringDB analysis depicts protein-protein interactions (PPI) between ExN1 marker NR4A2 TF (circled in blue) and MDD-associated TFs significantly enriched (q<0.05; Homer) in ExN1 DARs (circled in red) (interaction evidence>0.7 (high); direct interactions: FOS, ASCL1, FOSB and indirect: EGR1). E-F. Module scores computed using E. promoter-accessibility and F. snRNA-seq integrated gene-expression for GM and WM specific microglia marker genes. Boxplot shows significant differences in module scores between snATAC-seq microglial clusters (Mic1 and Mic2; Wilcoxon, p<0.05). G. 10x Visium spatial gene-expression in male and female DLPFC sections colored by the label-transfer prediction scores from snATAC-seq microglial clusters, Mic1 (top) and Mic2 (bottom), using promoter-accessibility (left) and snRNA-seq integrated gene-expression (right) H-I. Module scores computed using H. promoter accessibility and I. snRNA-seq integrated gene-expression for MDD-associated significantly down-regulated and up-regulated genes in cortical GM depression disease-associated microglia (depDAM)75. Boxplot shows significant differences in module scores between snATAC-seq microglial clusters (Mic1 and Mic2; Wilcoxon, p<0.05).

**Extended Figure 7:**
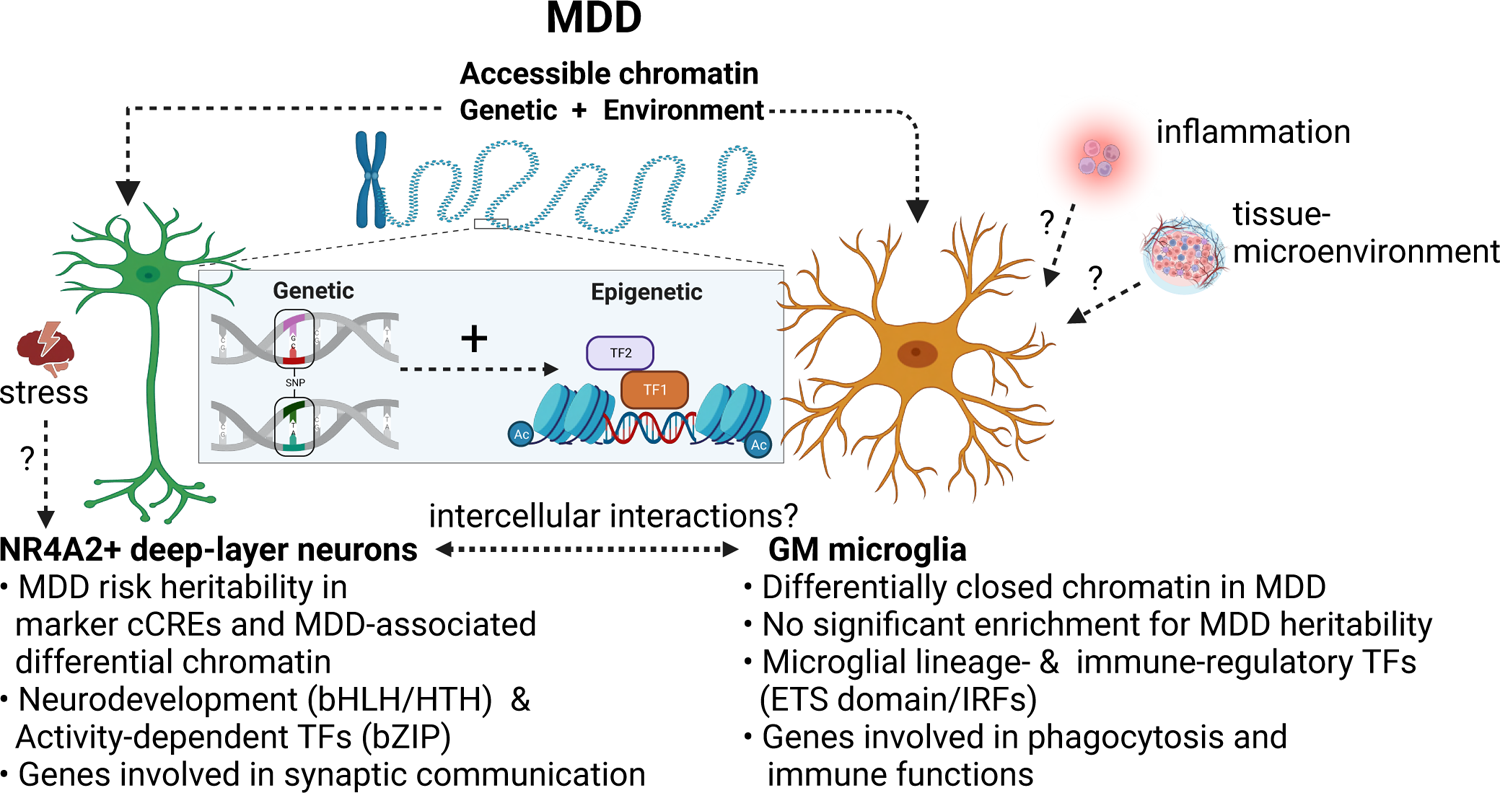
Proposed model for MDD associated regulatory mechanisms in deep-layer excitatory neurons (ExN1) and grey matter microglia (Mic2) through which genetic and epigenetic factors may mediate disease pathology.

**Supplementary Figure 1:**
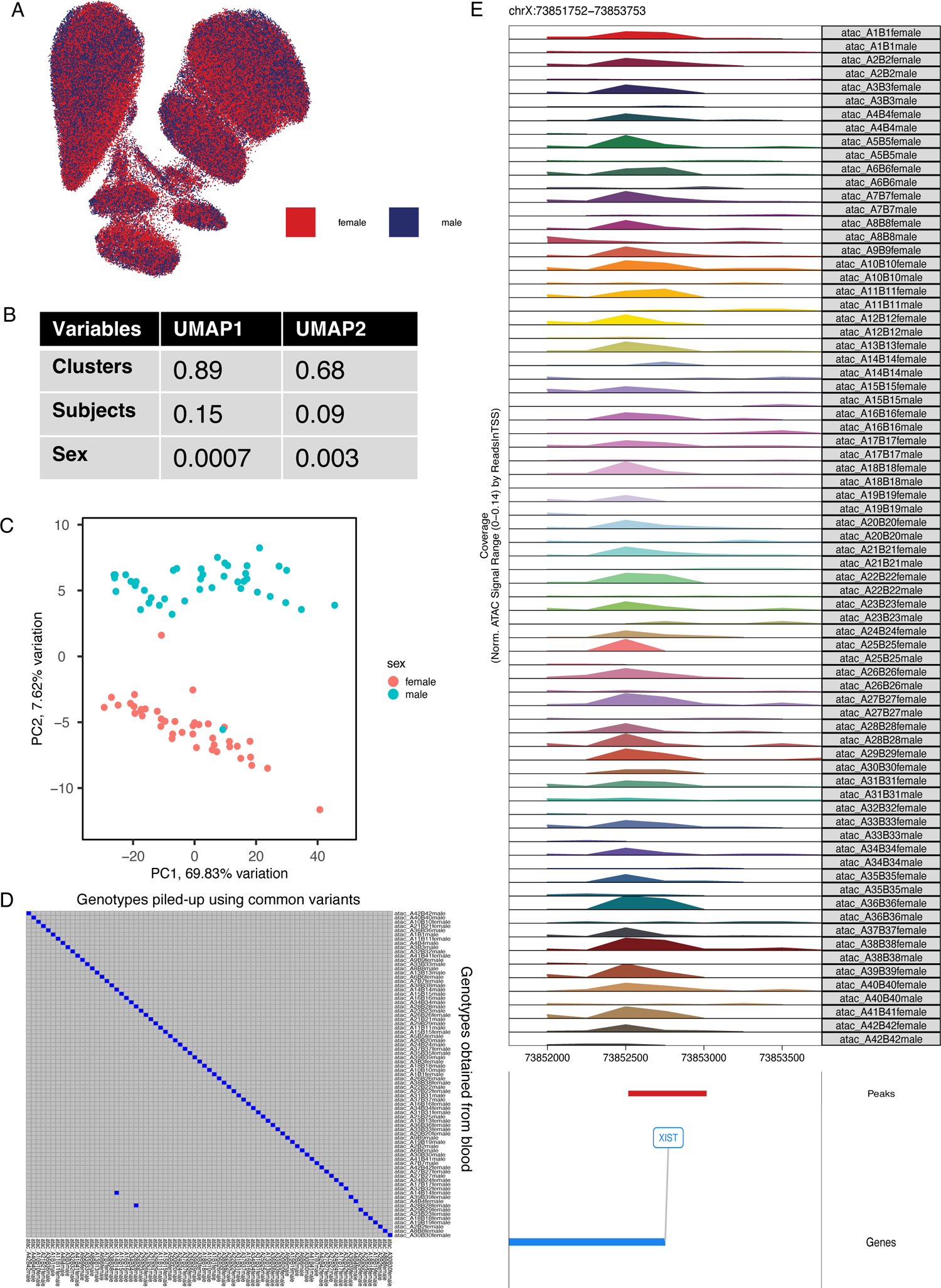
Demultiplexing of pooled male and female subjects. A. UMAP plot based on chromatin accessibility in 500bp genomic bins and colored by sex. B. The table shows R2 values reflecting cluster, individual and sex-specific variation explained by UMAP1 and UMAP2 embeddings. C. PCA of top 1% most variable cCREs identified using pseudo-bulked counts per subject and colored by the sex identified for each demultiplexed subject. D. Best match between genotypes of demultiplexed subjects piled-up using 1000 Genomes based common variants and genotypes obtained from the blood of these individuals. E. Pseudo-bulk chromatin accessibility (normalized by reads in TSS) at *XIST* gene in each of the demultiplexed subject.

**Supplementary Figure 2:**
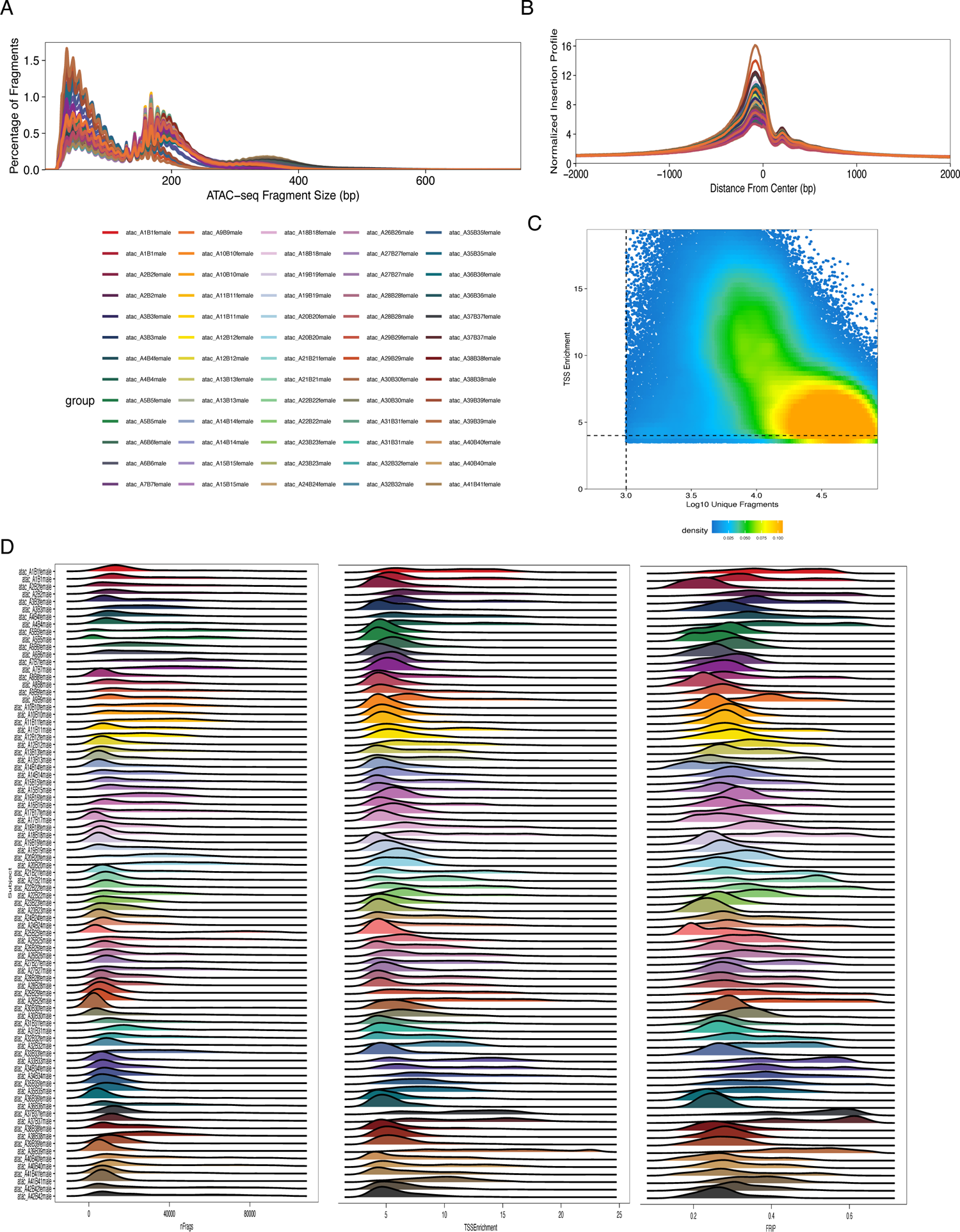
Quality-control of snATAC-seq. A. snATAC-seq fragment-size distribution in each subject. B. Normalized Tn5 insertions profiles aggregated across TSS in each subject. C. Density scatter plot depicting relationship between TSS Enrichment and log(10) unique fragments across 201456 cells. D. Ridge plots showing distribution of total number of fragments (nFrags), TSSEnrichment, and fraction reads in peaks (FRIP) in each of the demultiplexed subjects.

**Supplementary Fig 3:**
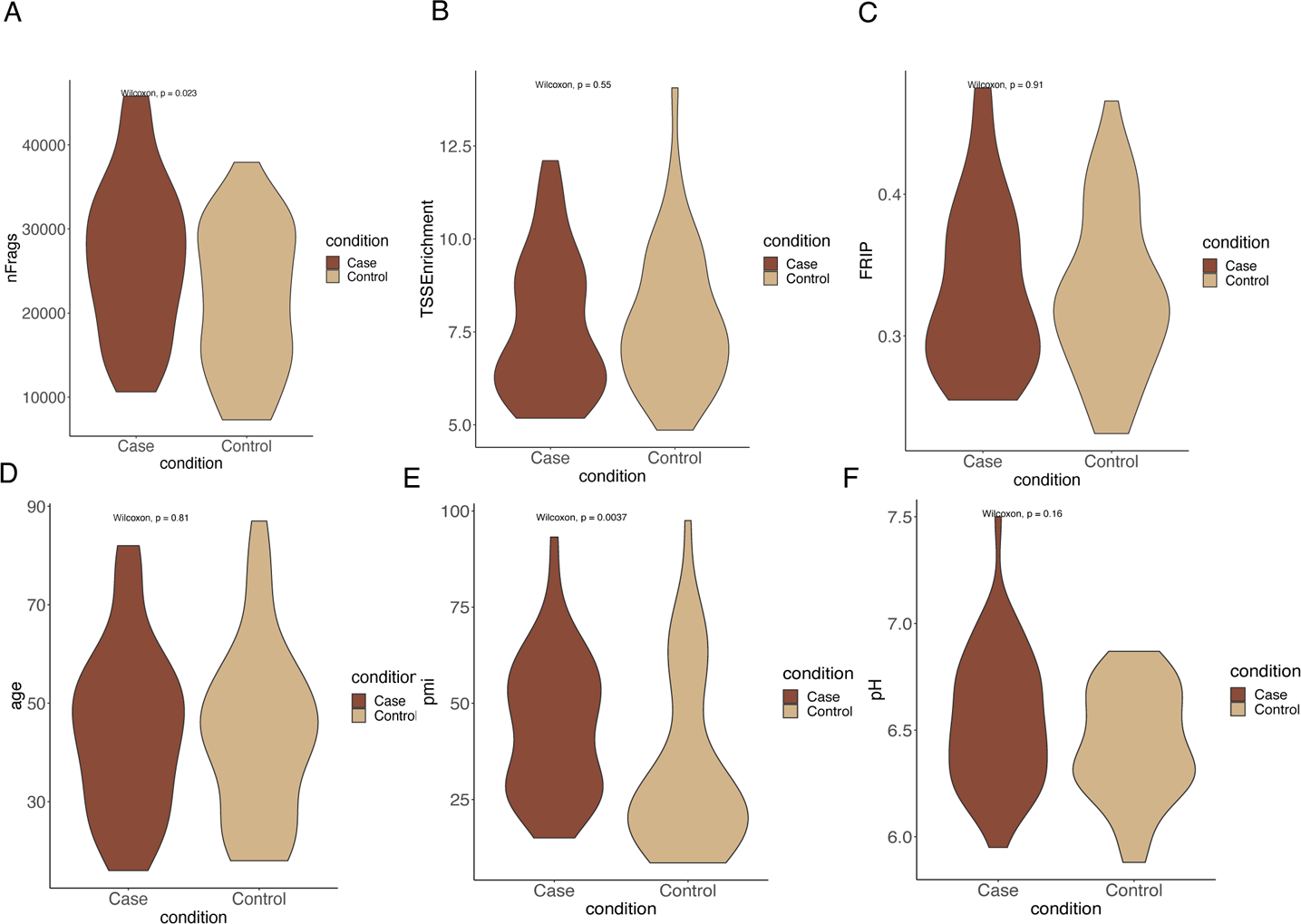
Quality-control of snATAC-seq. Comparison of following mean variable values across cells in MDD versus control subjects (Wilcoxon test) A. no. of fragments B. TSSEnrichment. C. fraction reads in peaks (FRIP). D. Age. E. post-mortem interval (PMI). F. pH of the brain tissue.

**Supplementary Fig 4:**
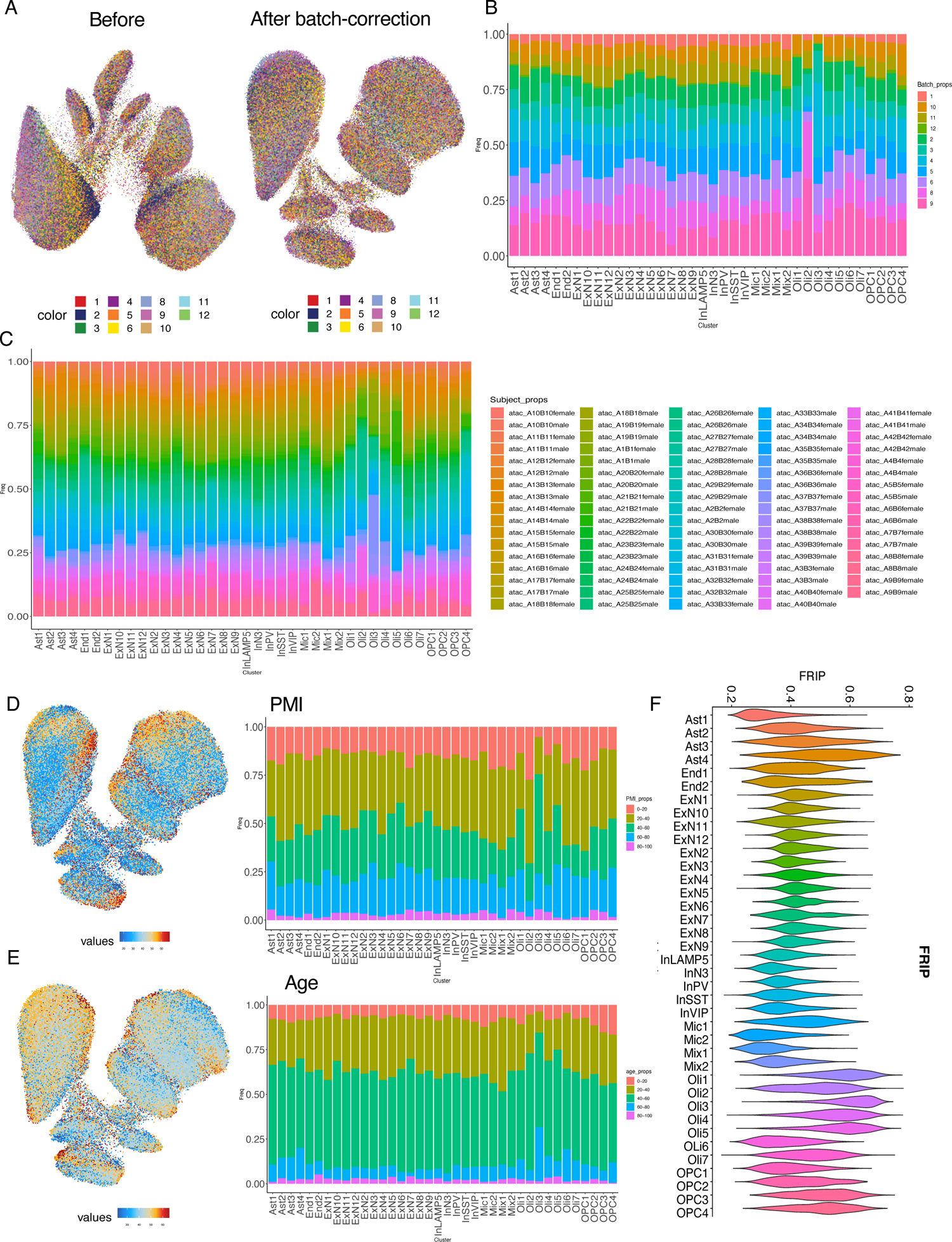
Quality-control of snATAC-seq. A. UMAP plots before (left) and after (right) batch-correction colored by batches. B. Bar plots showing proportions of cells contributed by each batch in each cluster. C. Bar plots shows proportions of cells contributed by each subject in each cluster. D. UMAP plot colored by PMI (left) and Bar plots (right) showing proportions of cells contributed by PMI groups in each cluster. E. UMAP plot colored by age of subjects (left) and Bar plots (right) showing proportions of cells contributed by each age group in each cluster. F. Violin plots depicting distribution of FRIP across individual cells in each cluster.

**Supplementary Figure 5:**
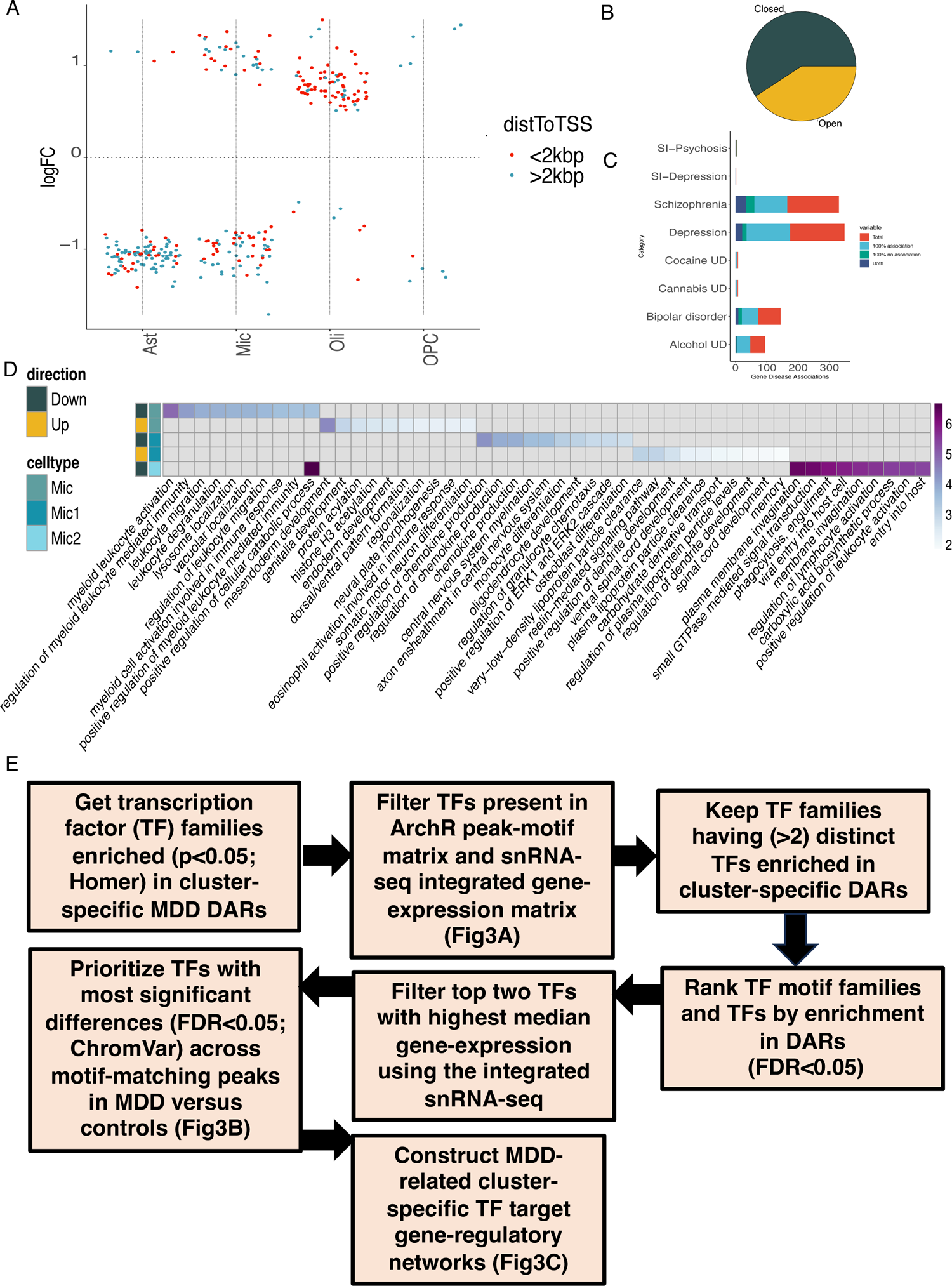
Differential accessibility in MDD. A. Differential accessibility in MDD versus control subjects in each cell-type (Limma-voom; FDR < 0.05 & abs(LogFC) > log2(1.1)) split by the direction of effect in MDD versus controls and colored by the distance to the nearest TSS. B. B. Pie chart depicts the total number of differentially accessible regions across cell-type DARs that are differentially more (open; n=121) or less accessible (closed; n=176) in MDD versus controls subjects. C. PsyGeNet analysis for psychiatric disorders shows the total number of gene-disease associations for genes linked to cell-type DARs (r>0.45). D. Heatmap (colored by –log10(p-values)) shows GO pathways (Top 10; FDR<0.2) enriched in genes linked (r>0.4) to differentially less (down) or more accessible (up) MDD DARs (FDR<5%) in broad microglia cell-type (Mic) and microglial clusters (Mic1, Mic2). E. Flowchart overview of TF prioritization workflow and construction of target gene-regulatory networks in MDD.

**Supplementary Figure 6:**
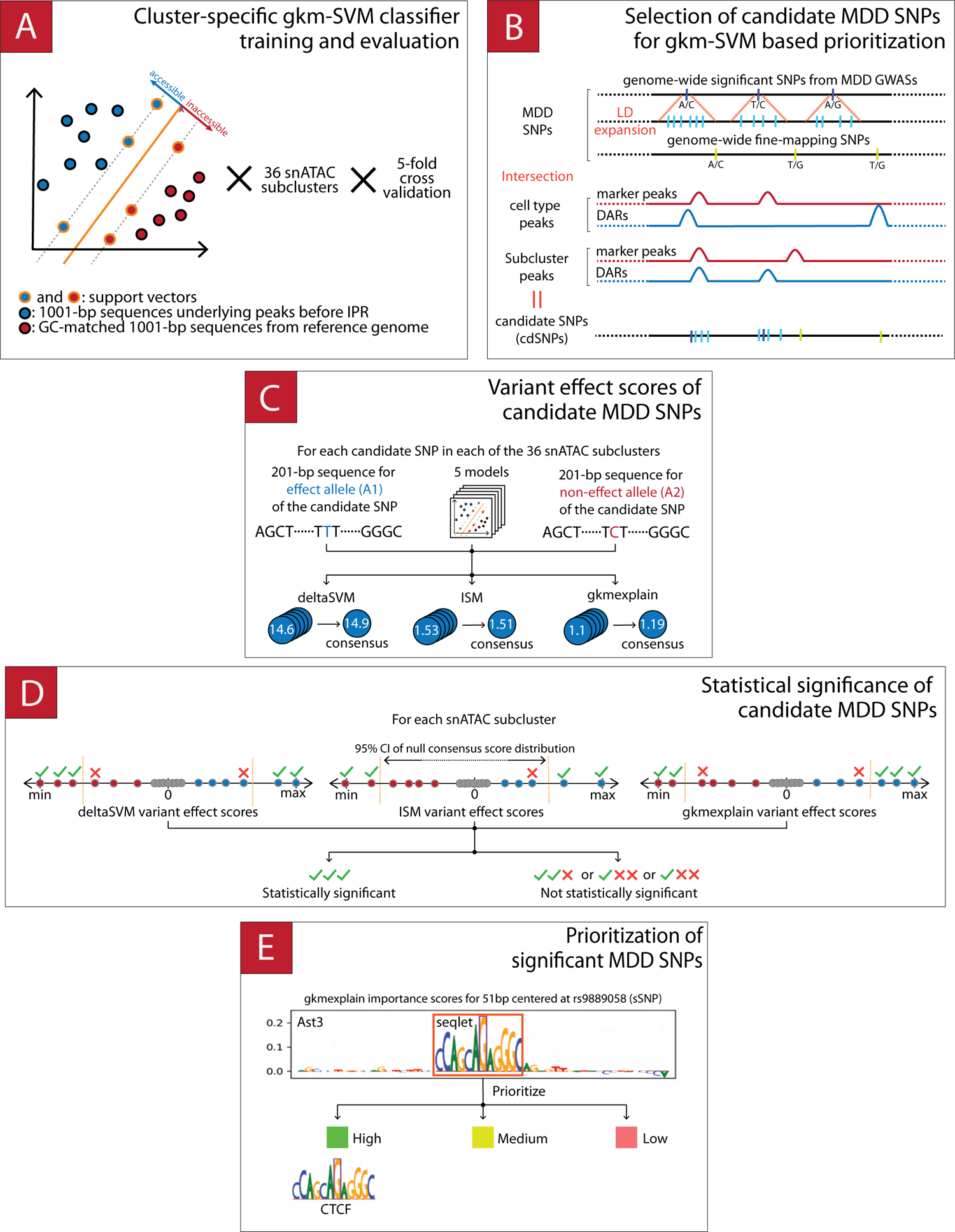
Schematic describing gkm-SVM based variant-scoring workflow employed for each snATAC cluster. A. Cluster-specific gkm-SVM classifiers were trained to predict cCRE status of input 1001-bp genome sequences via 5-fold cross validation. B. MDD SNPs located within 1001-bp marker cCREs and DARs are pooled to obtain candidate SNP set per cluster (cdSNPs). C. gkm-SVM-based cdSNP variant effect scores were computed by running ISM, deltaSVM and gkmexplain on 201-bp allelic sequences of cdSNPs. D. Statistical significance of cdSNPs were assessed with respect to score-specific null t-distributions fitted to null variant effect scores computed on shuffled (di-nucleotide preserved) 201-bp allelic sequences. For each score type and cluster, cdSNPs whose variant effect scores unanimously lie outside of 95%CI of respective null distributions were coined as statistically significant (sSNP). E. sSNPs were further partitioned into three groups according to the prominence and magnitude of their allelic impact to their immediate (seqlet) as well as 51-bp neighborhood. Additionally, seqlets of sSNPs having high or medium importance were interrogated for potential TFBS matches in CIS-BP 2.0 (TOMTOM, q-value < 0.1). Please see supplementary methods for implementation specific details regarding the workflow, and supplementary results for reliability analysis of workflow components.

**Supplementary Figure 7:**
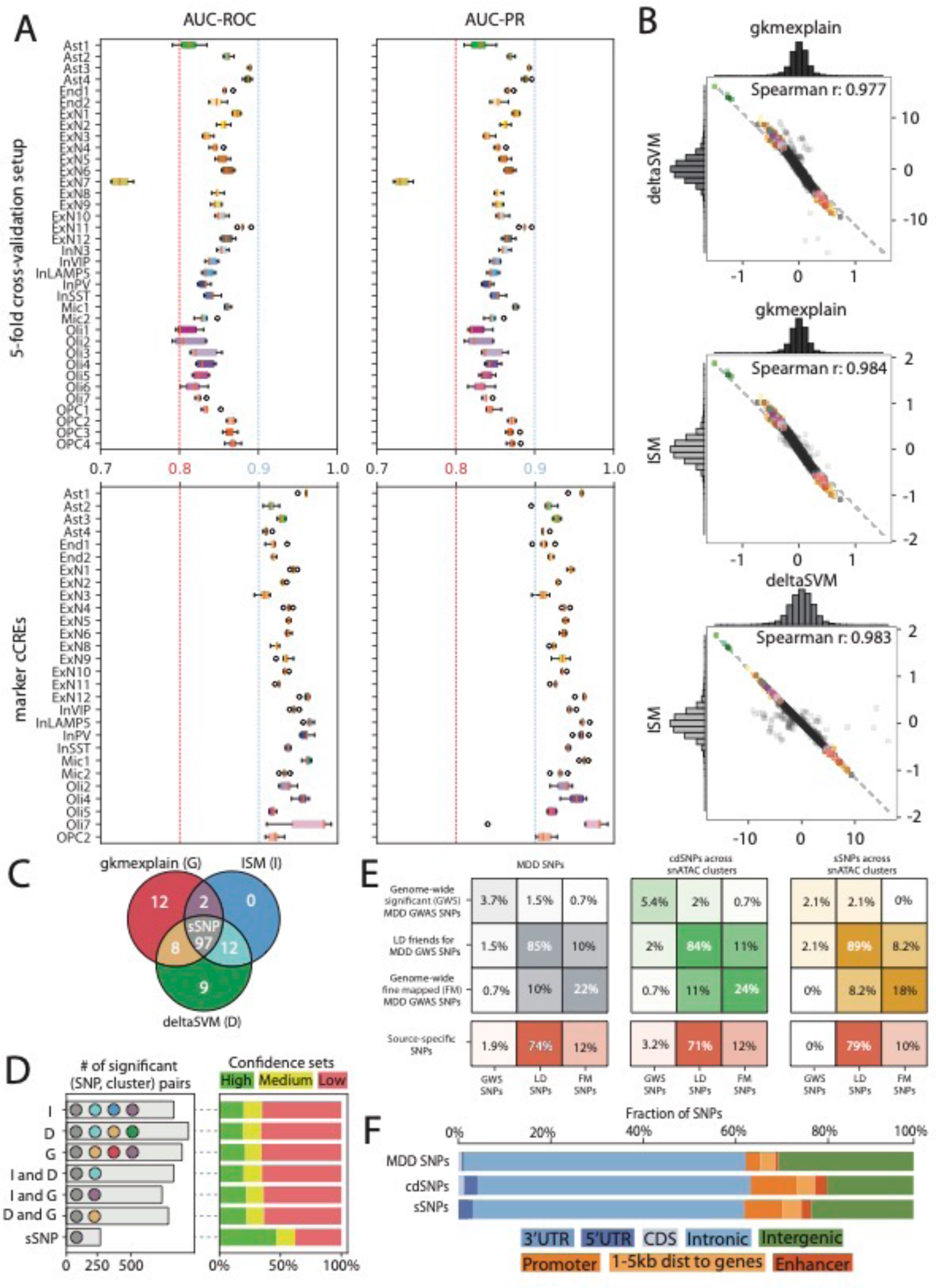
Cluster-specific SVM models for identification of MDD sSNPs. A. Performance comparison of cluster-specific gkm-SVM classifiers on cross validation test splits (top) and marker cCREs (bottom) with respect to AUC-ROC (left) and AUC-PR (right). B. Pairwise correlation of cdSNP variant effect scores across snATAC clusters. C. Statistical significance of cdSNPs according to ISM, deltaSVM and gkmexplain variant effect scores. D. Distribution of potentially significant cdSNPs into confidence sets. E. Composition of MDD SNP (left), cdSNP (middle) and sSNP (right). (Panel F) Genomic location based categorization of MDD SNPs (top), cdSNPs (middle) and sSNPs (bottom).

