## Supplementary tables for "Differential Chromatin Architecture and Risk Variants in Deep Layer Excitatory Neurons and Grey Matter Microglia Contribute to Major Depressive Disorder": Supplementary note.docx

### **Validation of demultiplexing approach adopted for snATAC-seq.**

We multiplexed nuclei extracted from a male and a female subject to reduce capture-cost and potential batch-effects between subjects and sexes (Supplementary Fig 1A-B; Supplementary Fig 4A). Principal component analysis (PCA) of top 1% most variable peaks clustered the demultiplexed subjects by sex (Supplementary Fig 1C; Supplementary method: Demultiplexing and assignment of sex). In addition, pseudo-bulk chromatin accessibility profiles at *XIST* distinguished demultiplexed donors by sex (Supplementary Fig 1E). One subject was removed from downstream analysis as we could not confirm the reported sex. Out of remaining 83 subjects used for downstream analysis, one subject (reported as male by the brain bank) clustered with the female sex in PCA (Supplementary Fig 1C); however, both snRNA-seq and genotyping consistently identified this subject as female, depicting robustness of our snATAC-seq based demultiplexing approach. We finally validated subject identities by assessing concordance between genotypes obtained from 1000 Genomes based common variants ^1^ and genotypes obtained from individual blood samples (Supplementary methods). Almost all the subjects matched correctly (Supplementary Fig 1D), except two female subjects that showed the best match with the multiplexed male subjects. Notably, both these female subjects clustered with the female sex in PCA and showed female-specific accessibility at *XIST* (Supplementary Fig 1). Taken together, we show that our sex-based multiplexing approach can allow for robust demultiplexing of subjects in snATAC-seq. Moreover, this method omits the need for sample genotyping and significantly reduces the cost of 10x single-cell capture and sequencing.

### **Functional MDD sSNPs identified using cluster-specific gapped-kmer SVM**

In total, 13,531 MDD associated SNPs were derived from three GWAS ^2^ ^3^ ^4^ consisting of LD expanded (86.3%), fine-mapped (22.3%), and genome-wide significant SNPs (3.7%). As expected, these SNP sets overlapped with each other, but each of them had their exclusive contributions to MDD associated SNPs (75.4%, 1.9% and 12%, respectively; Supplementary Fig 7E). Across 35 snATAC clusters (excluding ExN7, Mix1 and Mix2) and corresponding broad cell-types, we observed that 8.7% of MDD associated SNPs were overlapping with 1001-bp marker cCREs or candidate DARs (cdSNPs). In turn, 8.5% of cdSNPs were qualified as sSNP for at least one cluster by gkmSVM modelling. cdSNPs were relatively more abundant in broad cell-type and cluster marker peaks (43.7% and 83.4%) compared to DARs (2% and 12.5%). Further, 71.1% and 80% of sSNPs were observed in cell-type and cluster marker peaks, respectively.

Genomic location-based characterization of MDD associated SNPs, cdSNPs and sSNPs revealed a consistent overlap with intronic (ranging from 64.9% to 69.7%; Supplementary Fig 7F) and intergenic regions (ranging from 19.1% to 29.3%). Moreover, we observed that a greater percentage of cdSNPs and sSNPs were in promoter regions (11.1% and 12.7%), compared to MDD associated SNPs (3.8%). Additionally, the percentage of cdSNPs and sSNPs overlapping with 5’UTR were slightly greater than that of 3’UTR (0.5% and 1% increase) unlike the case with MDD associated SNPs (1.3% decrease).

Overall, we observed that majority of sSNPs were detected in neurons (78%), specifically excitatory neurons (75%), compared to glial cell types (22%). Moreover, 97.9% of the sSNPs were exclusively identified in clusters of a single broad cell-type as opposed to the few SNPs shared between multiple cell-types (2.1%; Ast-Oli and ExN-Mic). Furthermore, 53.1% of sSNPs were exclusively disrupting accessibility in a single cluster.

In case of excitatory neuronal clusters, we found sSNPs mapping to specific cortical layers, where the highest percentage of cluster-exclusive sSNPs were observed in deep-layer neurons. Notably, our candidate cluster ExN1 had the highest percentage of cluster-exclusive SNPs (40%) among any other deep-layer clusters identified in our data. Only a handful sSNPs were observed to have an effect shared across all excitatory neuronal clusters (1.4%). On the other hand, in case of inhibitory neuronal clusters, the percentage of sSNPs (3%) were much lower than that of excitatory neuronal clusters.

In case of glial clusters, we identified a variety of sSNPs for astrocyte (6.1%), oligodendrocyte (7.2%), OPC (3.1%) and microglia (4.1%) clusters. Of note, rs9889058 in astrocytes had the highest observed consensus variant effect score across all clusters. This sSNP was located in astrocyte cell-type candidate DAR and showed the most significant variant-effect score in Ast3 among other astrocytic clusters. This MDD sSNP was found to disrupt a Zf family CTCF TF binding site and was linked to *GNAO1* gene through peak-to-gene linkages. Interestingly, Ast3 cluster also showed significant accessibility disruption in MDD individuals (followed by ExN1 and Mic2). Likewise, rs174561 sSNP was located in cluster-specific marker peaks of an astrocytic Ast4 and oligodendrocyte Oli1 clusters in promoter, exon, intron and 5’UTR regions of various *FADS1* transcripts, and was eQTL for *FADS1*, *FEN1*, *TMEM258*, *NXF1* as well as sQTL for *FADS3* gene.

### **Reliability assessment of cluster-specific SNP prioritization**

Here, we interrogate individual components of gkmSVM based workflow to assess whether statistical significance analyses of cdSNPs is reliable.

The trained classifiers for all clusters except ExN7 attained between 80% and 90% median AUC-ROC and AUC-PR computed on held out test folds of cluster-specific 5-fold cross validation schemes (Supplementary Fig 7A). Regardless of the performance of ExN7 models, this cluster was omitted from downstream analyses due to low signal-to-noise ratio. Additionally, for 28 clusters, we tested the trained models on datasets consisting of 1001bp sequences underlying cluster marker peak regions as well as GC content and chromosome matched non-peak sequences. We observed that the models had even better performance on marker peaks with respect to median AUC-ROC (ranging from 90.7% to 98.2%) and AUC-PR (ranging from 91% to 98.1%). Since most of the cdSNPs were in cluster marker peak regions, we concluded that predictive power of the trained models was suitable for conducting a reliable chromatin accessibility disruption analysis.

Next, we examined the agreement between consensus ISM, deltaSVM and gkmexplain variant effect scores. We observed that each pair of scores were significantly correlated with respect to cdSNP variant effect scores (Spearman’s 𝜌, min: 0.977, max: 0.984). Since none of the score type pairs had perfect correlation (Supplementary Fig 7B-C), we further investigated the cdSNPs which were not eligible to be sSNPs but were deemed statistically significant with respect to at least one score type. In line with the high observed correlations, number of such SNPs were much lower compared to sSNPs (Supplementary Fig 7D). Hence, incorporating all three score types in sSNP criteria makes our analysis more stringent and reduces false positive findings.

Finally, we investigated the percentage of high, medium and low confidence sets of sSNPs whose seqlets were successfully matched (q-value < 0.1) to TFBS via TOMTOM. We observed that a greater percentage of high and medium priority sSNPs matched to TFBS (47.2% and 45.2%; Supplementary Fig 7D), compared to low priority sSNPs (18.4%).

### **Supplementary Methods**

### Nuclei Extraction

Nuclei were extracted from frozen DLPFC tissue sections as previously described ^5^ with some modifications. Briefly, frozen brain sections were dounced for 5mins in the lysis buffer. The lysis suspension was homogeneously mixed by pipetting up and down 10-15 times following which only 80% of the lysis solution was mixed with 5ml of wash buffer to make diluted nuclei suspensions. The samples were mixed and passed through a 30-um cell strainer to remove cellular debris and washed twice in 10ml wash buffer. Next, Optiprep cushion was created followed by gradient centrifugation for 30mins at 10,000g and resuspended in 1ml wash buffer. Nuclei were counted with Hoescht 33342 (1:2000) three times (average was used) with Cell countess (Olumpys). Nuclei were then spun down at 500g for 5mins and diluted in 1xNB (10x Genomics) to obtain a final concentration of 3080 nuclei/ul. We increased the loading concentration by 20% to compensate for potential sources of nuclei loss as done previously ^5^ ^6^.

### Multiplexing and library construction

Equal volume of nuclei suspensions from a male and female subject (either from case or control) were pooled together to obtain a final concentration of 3080 nuclei/ul. The combined nuclei were loaded in one of the lanes of 10x microfluidic chip. Each microfluidic chip was loaded with at least four multiplexed libraries representing both cases and controls. Single-cell capture and library preparations were done using 10x Genomics Chromium Single Cell v1.1 reagents following the protocol outlined by the user guide [[https://www.10xgenomics.com/support/single-cell-atac/documentation/steps/library-prep/chromium-single-cell-atac-reagent-kits-user-guide-v-1-1-chemistry].](https://www.10xgenomics.com/support/single-cell-atac/documentation/steps/library-prep/chromium-single-cell-atac-reagent-kits-user-guide-v-1-1-chemistry%5d.) Cell Ranger atac v2.0 was used for alignment against the GRCh38 reference available on the 10x Genomics website (refdata-cellranger-arc-GRCh38-2020-A-2.0.0). The sequencing metrics are provided in Supplementary Table 1.

### Demultiplexing and assignment of sex

Male and female nuclei pooled 10x snATAC-seq libraries were demultiplexed at single-cell level using Vireo ^7^ into specified number of donors (n=2) based on 1000 Genomes based common variants ^1^ piled-up at single-cell level using CellSNP ^8^. Further, the genotyping data obtained from blood was lifted over from hg19 to hg38 using LiftoverVcf from picard. CellSNP-lite ^8^ was used to call variants from snATAC-seq demultiplexed subject bam files with the --minMAF flag set to 0.1 and the --minCOUNT flag set to 20. Vireo with --no-doublet flag option was used for donor deconvolution using genotypes obtained from blood and mismatches were assessed between the genotyping array data and snATAC-seq based demultiplexing approach. Finally, to determine sex of the demultiplexed donors, we used multiple approaches: a) PCA of top 1% most variable peaks using pseudo-bulked peak accessibility counts per subjects, b) psuedobulk accessibility profiles (normalized by reads in TSS) at sex-specific gene *XIST,* c) finally, to confirm sex at single-cell level, we trained a machine learning classifier on sex-specific chromatin accessibility data generated previously (data not shown). Overall, these results agreed with pseuodbulk accessibility profiles of each subject identified by PCA and at sex-specific *XIST* gene.

### Genotyping Array Processing

GenomeStudio was used to call genotypes from raw idat files and the genotypes were exported from genome studio according to Illumina’s Plus/Minus convention. After filtering out SNPs with a call rate less than 95%, MAF < 0.01, or Hardy-Weinberg equilibrium p-value less than 0.000001, the genotypes were phased with SHAPEIT v2 using 15 burn in iterations, 8 pruning iterations, 40 main iterations, 200 states, and a random seed of 1. Imputation was then performed on 3mb segments of each chromosome using IMPUTE2 v2.3.2. For both phasing and imputation, the GRCh37 1000 Genomes phase 3 dataset was used as the reference. After imputation, SNPs with MAF < 0.01, Hardy-Weinberg equilibrium p-value less than 0.000001, or INFO score < 0.8 were removed.

### Quality Control of Genotyping Array Samples

Sex mismatches were identified in the dataset using PLINK 1.9's --check-sex option to calculate the X chromosome inbreeding coefficient. Subjects annotated as male with X chromosome inbreeding coefficients less than 0.8, or annotated as female with inbreeding coefficients greater than 0.2 were deemed problematic. Potential duplications of samples as well as possible cross contamination of samples was examined using identity by descent calculated by PLINK 1.9's --genome option, after LD pruning. Samples that were mismatched in both the snATAC-seq and snRNA-seq datasets were removed from the genotyping array dataset. Samples with call rates less than 98% were also removed.

### PsychEncode data

PsychEncode DLPFC neuronal and non-neuronal (NeuN+/-) HiC loops ^11^ and NeuN+/- H3K27ac and H3K4me3 histone modification peaks were obtained ^12^. For integrating NeuN+/- histone peaks in MDD sSNP accessibility tracks, we used per-subject DLPFC histone peaks (bed format) from the PsychEncode EpiMap database [[https://psychencode.synapse.org/Explore/Studies].](https://psychencode.synapse.org/Explore/Studies%5d.) and generated a merged non-overlapping peak-set across all subjects using GenomicRanges reduce() function. Peaks were converted to hg38 using kentutils liftOver command line with hg19ToHg38.over.chain.gz UCSC liftover chain. For assessing overlaps with DARs, marker cCREs, and peak-to-gene linkages, we used consolidated histone modification peaks from cortical cells ^12^.

### MDD GWAS fine-mapping

Genome-wide fine-mapping of MDD GWAS ^2^ with SparsePro ^13^ was performed on summary statistics and reference LD panel based on White British population of UK Biobank ^14^. Briefly, SparsePro utilizes variational inference to (a) aggregate correlated variants having similar association with a trait into 𝐾 sparse effect sets; (b) infer set-level causal status, effect sizes and configuration. Moreover, effect sizes as well as causal status and configuration of effect sets were used to compute variant-level posterior inclusion probability (PIP) and effect size estimates. Fine-mapping inputs were prepared as follows: GWAS variants were first binned into sliding windows (i.e., loci) of 3-Mbp with neighbouring windows having a 2-Mbp overlap. In turn, loci-level summary statistics were harmonized with LD matrix by inverting effect size and alleles of mismatching SNPs. SparsePro was run with 𝐾=5 sparse effect sets on each 3-Mbp loci. To mitigate boundary effects, estimates for the variants located within 1-Mbp windows centered at each loci were retained. Moreover, 95% credible set was computed on genome-wide variant-level PIP estimates. Next, sparse effect sets inferred at each loci were filtered to retain those including at least one variant qualified for the credible set. Across 22 autosomal chromosomes, 213 independent effect sets comprising of 3,013 variants in LD were identified to be likely causal for MDD across the genome. The nearest genes and e/sQTLs associated with these variants are provided (Supplementary Table 14).

### Cluster-specific accessibility disruption analyses by MDD associated genetic variants.

We adapted gkmSVM variant-scoring workflow ^15^ (Supplementary figure 6) consists of the following steps: (a) gkmSVM training and evaluation; (b) gkmSVM-based consensus variant effect scores of cdSNPs; (c) Statistical significance of cdSNPs; and (d) High, medium and low confidence subsets of sSNPs.

gkmSVM training and evaluation

For each of the 35 snATAC clusters (excluding Mix1, Mix2, and ExN7), we adopted a 5-fold cross validation setup and trained gkmSVM models to predict whether 1001bp genomic sequences are from cluster-specific cCREs (accessible, positive examples) or randomly sampled non-peak regions (inaccessible, negative examples) using default parameter setting defined as follows: gapped k-mer kernel with center weighting (wgkm, t = 4); a word length of 11 (l = 11); 7 informative columns in each word (k = 7); the maximum allowed number of mismatches to consider (d = 3); the initial value of the exponential decay function used for center weighting (M = 50), half-life parameter of 50 positions for exponential decay (H = 50); the regularization penalty parameter of SVM (c = 1); and precision parameter (e = 0.001).

The datasets used for each cluster consisted of equally numerous positive and negative examples with matching chromosome and GC content distributions. To obtain positive example set per cluster, we applied iterative peak removal (based on MACS2 p-value) on 501bp peaks called for each cluster; extended resulting peaks by 250bp at both directions; extracted underlying sequences from the reference genome (hg38); and discarded the sequences containing an ambiguous base (“N” or “n”). In case of negative examples, we first generated a cluster-independent set of candidate negative examples by tiling hg38 reference genome 1001bp at a time with a stride of 50bp while discarding those having an ambiguous base. Moreover, individually for each cluster, we further removed the sequences intersecting with 1001-bp peak regions. Since the number of negative examples were much greater than those of positive examples in each cluster, we subsampled the former to match the size of the latter. For this goal, we first partitioned each positive example set into 20 equally populated bins according to the GC content distribution percentiles defined on the same set; and assigned the candidate negative examples according to the resulting cluster-specific bins. Next, we initialized an empty dataset per cluster and added pairs of positive and negative examples, one at a time, until the positive example set of the corresponding cluster is exhausted. At each step, we paired one randomly sampled positive example with one randomly sampled candidate negative example originating from the same chromosome and GC bin, where sampling was done without replacement. Resulting cluster-specific datasets were partitioned to cross-validation folds as follows: Fold 1 consisted of chromosomes 3, 4, 12 and 13; Fold 2 consisted of chromosomes 5, 6, 14 and 15; Fold 3 consisted of chromosomes 7, 8, 11, 16 and 17; Fold 4 consisted of chromosomes 2, 9, 10, 18 and 19; and Fold 5 consisted of chromosomes 1, 20, 21, 22, X and Y.

At each iteration of the cross-validation setup, gkmtrain function ^16^ was run with default parameter setting to train gkmSVM model on 4 folds; and gkmpredict^16^ function was run on sequences in the remaining (test) fold to obtain unnormalized decision scores whose magnitude quantified model confidence and sign reflected whether the sequences are accessible (i.e., positive). To make model training computationally feasible, we sorted positive and negative example pairs in each training set (i.e., 4 folds) with respect to the ascending order of MACS2 p-value of positive examples and, trained the models on the top 60,000 pairs (120,000 examples). End2, InN3 and Mic2 clusters did not have 60,000 positive examples in their training sets and all available pairs of examples were used for model training. Finally, the trained models (175 in total) were evaluated with respect to AUC-ROC and AUC-PR metrics (scikit-learn package ^17^) calculated according to test fold predictions (Supplementary Fig 7A).

gkmSVM-based consensus variant effect scores of cdSNPs

Here, we utilized the gkmSVM models to interrogate whether cdSNPs have an allelic impact on chromatin accessibility in the corresponding cluster. First, we extracted 201bp sequences centered at each cdSNP in each cluster from the reference genome (hg38); and generated two sequences per SNP by mutating the middle position (101st position) to the effect (A1) and non-effect (A2) alleles of the SNP, respectively. We employed three complementary approaches on 201-bp cdSNP sequences — in silico mutagenesis (ISM), deltaSVM ^18^ and gkmexplain ^19^ — to compute cluster-specific variant effect scores as follows:

To obtain *ISM* variant effect score per cdSNP in each cluster, we first executed gkmpredict function to predict whether 201bp allelic sequences of cdSNPs available for the corresponding cluster are accessible. Next, ISM variant effect score was computed by subtracting non-effect allele score from that of the effect allele resulting from the same model. Finally, consensus ISM variant effect scores for each cdSNP were calculated by averaging variant effect scores across the models trained for the corresponding cluster.

In the case of *deltaSVM*, we first generated and predicted all possible non-redundant 11-mers with the trained models via gkmpredict. Resulting decision scores served as a model-specific basis for deltaSVM score calculation. Next, for each trained model in each cluster, deltaSVM.pl script was run on 201-bp allelic sequences cdSNPs as well as 11-mer decision scores. Finally, we averaged resulting variant effect scores across models to get consensus deltaSVM variant effect scores per cdSNP.

Unlike ISM and deltaSVM which report magnitude and direction of change in gkmSVM predictions between sequence pairs, *gkmexplain* measures contribution of the nucleotides in each input sequence on gkmSVM predictions in terms of base-resolution importance scores ^19^. For each pair of cluster-specific cdSNPs and gkmSVM models, we first executed gkmexplain script on 201bp allelic sequences in importance score mode (-m 0) to obtain 201-by-4 gkmexplain allelic importance scores. Resulting matrices capture importance of base identities (i.e., A, G, C and T) available at each position (i.e., 201-bp) on model predictions and, thus only the base identities present at each position are non-zero. Next, we computed 201-by-4 consensus importance score matrices for each allele of cdSNPs as the mean of importance score matrices across models. Moreover, we extracted the consensus importance scores corresponding to the 51bp window centered at the cdSNP (i.e., from 75th to 126th position) and computed the respective sums of 51-by-4 consensus importance score matrices of both alleles. Finally, for each cdSNP in each cluster, consensus gkmexplain variant effect scores were computed by subtracting non-effect allele sum from that of effect allele.

Statistical significance of cdSNPs

To assess whether consensus variant effect scores of cdSNPs were statistically significant, we generated 10 randomly shuffled 201bp non-effect allele sequences (hereafter, null sequences) using fasta-dinucleotide-shuffle function of MEME Suite ^20^, followed by in silico mutation of the middle position of resulting sequences (i.e., 101st position) to both alleles of each cdSNP. Moreover, we computed null consensus variant effect scores for each score type (i.e., ISM, deltaSVM and gkmexplain) by repeating the previous step pairs of null allelic sequences resulting from the same random shuffle. Additionally, we fitted normal and t distributions using SciPy ^21^ to null consensus variant effect scores available for each score type per cluster. In line with the previous work ^15^ ^22^, t-distribution was found to be a better fit to empirical null consensus variant effect score distributions based on KS test. Finally, we denoted cdSNPs as sSNP within context of a cluster if and only if consensus variant effect scores for all score types reside outside of 95% confidence interval (CI) of respective null variant effect score distributions of the corresponding cluster.

High, medium and low confidence sets of sSNPs

To further refine sSNPs, we identified high, medium and low confidence sets reflecting the quantitative support for SNP-driven accessibility disruption in its immediate local neighborhood (<201bp). First, we determined the *active allele* for each sSNP which had a larger sum of non-negative consensus gkmexplain importance scores within the 51bp region centered at the SNP compared to the other allele. Next, we constructed cluster-specific and position-independent null importance score distributions by pooling all positive scores in 201-by-4 null consensus gkmexplain score matrices computed for cdSNPs of the corresponding cluster. Moreover, we extracted the subsequences around sSNPs (hereafter, *seqlets*) whose base-resolution consensus importance scores are significantly elevated compared to the position-independent null consensus importance distribution as follows: Starting from the position of sSNP, we extended an empty sequence towards both directions by iteratively adding the flanking nucleotides in an alternating fashion. If two consecutive positions in a direction had consensus importance scores lower than 97.5th percentile of the position-independent null distribution of the corresponding cluster, we stopped extending towards that direction. Furthermore, if the resulting seqlet was smaller than 7bp upon termination of the procedure, it was alternatingly extended towards both directions to reach this minimum length.

Next, we computed *prominence* and *magnitude scores* which depend on *seqlet score* and *seqlet signal-to-noise ratio (SNR)* scores based on seqlets for sSNPs in each cluster. Seqlet score for a given allele of an sSNP, is defined as the sum of non-negative consensus importance scores which overlap with seqlet of the SNP. Seqlet SNR for each allele of an sSNP is given by the ratio of seqlet score and the sum of non-negative values in 201-by-4 consensus importance score matrix. In turn, magnitude and prominence scores are given by the difference between non-effect and effect allele seqlet scores and seqlet SNRs, respectively. Magnitude score attempts to quantify the magnitude of chromatin accessibility disruption with respect to the change from non-effect to effect allele within seqlet region. On the other hand, prominence score attempts to capture how prominent the chromatin accessibility disruption due to the SNP is within its 201bp neighborhood. Furthermore, we sought to assess the statistical significance of the prominence and magnitude scores. For this goal, we first computed null prominence and magnitude scores by repeating the above-mentioned procedure on null sequences of cdSNPs in all clusters. Next, we calculated prominence and magnitude score p-values for each sSNP using cluster-specific empirical null distributions constructed on null magnitude and prominence scores as well as their additive inverse.

Finally, we partitioned sSNPs in each cluster to high, medium and low confidence sets according to the prominence and magnitude p-value based rule set defined here ^15^. sSNPs having prominence score p-value less than 0.05 were assigned to high confidence set. These sSNPs disrupt chromatin accessibility of their seqlet regions which were substantially more accessible compared to other regions within 201bp window centered at the SNP. Next, the sSNPs which did not qualify for high confidence set were assigned medium confidence if their magnitude score p-value were less than 0.05 or their prominence score p-value were less than 0.1. These SNPs substantially disrupt chromatin accessibility of their seqlet region but the quantitative support for disruption is moderate. Remaining SNPs were grouped under low confidence set. Additionally, we ran TOMTOM ^23^ (via memes R package ^24^) on seqlets of high and medium confidence sSNPs and identified significant matches (q-value < 0.1) to TFBS motifs (Homo Sapiens) available in CISBP-2.0 ^25^ database.

### DLPFC eQTL and sQTLs

A total of 8 eQTL (RNA-seq) and sQTL (LeafCutter ^26^) summary statistics corresponding to uniformly performed analysis on DLPFC tissue of individuals in BrainSeq ^27^, CommonMind ^28^, ROSMAP ^29^ and GTEx ^30^ studies were downloaded from eQTL Catalogue ^31^. We identified eQTLs and sQTLs with FDR<0.05 and used them for our downstream analyses.

### Genomic location-based characterization of SNPs

MDD associated SNPs, cdSNPs and sSNPs were characterized with respect to their genomic locations using several approaches. First, we downloaded human gene annotations (comrepensive set, hg38 build) from GENCODE v43 ^32^ and identified the genes nearest to each SNP. Moreover, we used annotatr R package ^33^ which outsources genic annotation data from org.Hs.eg.db and TxDb.Hsapiens.UCSC.hg38.knownGene R packages, to obtain fine-grained functional information regarding the genomic regions containing the SNPs. Specifically, we focused on the following annotation categories provided in decreasing order of priority: coding sequence (CDS), 5’UTR, 3’UTR, Exon, Enhancer (FANTOM5 ^34^ ^35^), Promoter (less than 1KBp distance to TSS), Intron, 1-to-5Kbp to TSS and Intergenic. Additionally, we determined DLPFC eQTL and sQTL hits (eQTL Catalogue, FDR < 0.05) within MDD associated SNP, cdSNP and sSNP sets.

### Characterizing MDD sSNPs-associated risk genes

For visualizing all the nearest or linked risk-genes to MDD sSNPs, we computed “full String network” with organism=”Homo Sapiens” and minimum required interaction score = 0.4 and maximum number of interactions (1^st^ shell= 0, 2^nd^ shell= 0). We then evaluated Monarch ^36^ based human phenotypes enriched in the network which resulted in the expected phenotypes, such as “depressive symptoms” and “wellbeing measurement.” Genes associated with sSNPs were input to enrichGO(minGSSize=5, pAdjustMethod="BH", pvalueCutoff=0.05, ont="ALL") using R package, clusterProfiler ^37^. Molecular pathways and human phenotypes enriched for sSNP target genes are provided (Extended figure 4, Supplementary Table 13).
